# CAMP: A modular metagenomics analysis system for integrated multi-step data exploration

**DOI:** 10.1101/2023.04.09.536171

**Authors:** Lauren Mak, Braden Tierney, Wei Wei, Cynthia Ronkowski, Rodolfo Brizola Toscan, Berk Turhan, Michael Toomey, Juan Sebastian Andrade Martinez, Chenlian Fu, Alexander G Lucaci, Arthur Henrique Barrios Solano, João Carlos Setubal, James R Henriksen, Sam Zimmerman, Malika Kopbayeva, Anna Noyvert, Zana Iwan, Shraman Kar, Nikita Nakazawa, Dmitry Meleshko, Dmytro Horyslavets, Valeriia Kantsypa, Alina Frolova, Andre Kahles, David Danko, Eran Elhaik, Pawel Labaj, Serghei Mangul, The International MetaSUB Consortium, Christopher E. Mason, Iman Hajirasouliha

## Abstract

**Motivation:** Computational analysis of large-scale metagenomics sequencing datasets have proven to be both incredibly valuable for extracting isolate-level taxonomic, and functional insights from complex microbial communities. However, due to an ever-expanding ecosystem of metagenomics-specific methods and file-formats, designing seamless and scalable end-to-end workflows, and exploring the massive amounts of output data have become studies unto themselves. One-click bioinformatics pipelines have helped to organize these tools into targeted workflows, but they suffer from general compatibility and maintainability issues, and preclude replication.

**Methods:** To address the gap in easily extensible yet robustly distributable metagenomics workflows, we have developed a module-based metagenomics analysis system “Core Analysis Modular Pipeline” (CAMP), written in Snakemake, a popular workflow management system, along with a standardized module and working directory architecture. Each module can be run independently or conjointly with a series of others to produce the target data format (e.g. short-read preprocessing alone, or short-read preprocessing followed by *de novo* assembly), and outputs aggregated summary statistics reports and semi-guided Jupyter notebook-based visualizations.

**Results:** We have applied CAMP to a set of ten metagenomics samples to demonstrate how a modular analysis system with built-in data visualization at intermediate steps facilitates rich and seamless inter-communication between output data from different analytic purposes.

**Availability:** The CAMP ecosystem (module template and analysis modules) can be found https://github.com/Meta-CAMP.

## Introduction

Metagenomics - the study of microbial communities recovered directly from environmental samples using genomic methods - has emerged over the last two decades to become an important field in bioinformatics. This culture-independent method of investigating microbial communities in their natural contexts has revealed invaluable insights that have greatly enhanced our understanding of the natural world [Almeida et al., 2019, Danko et al., 2021, Kadosh et al., 2020, Brito et al., 2019, Sierra et al., 2022]. Despite a vast body of literature, a rapidly expanding toolkit, and several established protocols, metagenomics still has a long way to go due to the complex nature of the microbial world and the innate challenges of analysis workflows combined. Since a microbial community typically consists of a complex mixtures of dozens to thousands of species spanning both known and unexplored branches of the tree of life [Bahram et al., 2021, Hug et al., 2016], precise and reliable workflows are required to identify these species and decipher their functional roles. In a world today where global-scale sequencing efforts such as Human Microbiome Project (HMP) [Gevers et al., 2012, Huttenhower et al., 2012] and International Metagenomics and Metadesign of Subways and Urban Biomes (MetaSUB) [Danko et al., 2021] are underway, we have the opportunity to work with extremely rich, deep, and high-dimensional sequencing datasets. There is a field-wide need to develop workflow systems that are straightforward to test, maintain, reuse to generate reproducible results [Djaffardjy et al., 2023].

A typical metagenomic workflow consists of multiple *in silico* steps following sample collection and sequencing to resolve the taxonomic and/or functional composition of the originating community [Quince et al., 2017]. Over the past decades, a myriad of tools have been developed and are continually developed and updated with the goal of supporting the various components of these analytic goals. Despite the presence of a vast array of open-source tools available, many of them are not accessible online, easy to install, or even installable at all [Mangul et al., 2019]. A widespread problem in bioinformatics is the short life span of tools. Kern et al. [2020] reported that of the 2,396 web tools they surveyed in a 133-day test frame, only 31% of tools always worked, 48.4% occasionally and 20.6% never worked.

This issue of fragility of tools cascaded when these tools are organized into workflows, resulting in many dependent points of potential failure at both installation and run times. Although open-source package management systems such as Conda, containerization systems such as Docker and Singularity, and workflow management systems such as Nextflow and Snakemake, have partially addressed these challenges, dependency conflicts and operating incompatibilities are still major problems when multiple tools need to co-exist in the same environment [Merkel, 2014, Kurtzer, 2018]. Besides, workflows that solely rely on containerization for environment management are not widely usable on high-performance compute clusters with root-access restrictions.

To streamline the complex process of decision-making, especially for users without extensive computational backgrounds, all-in-one pipelines such as ATLAS [Kieser et al., 2019], FinnLab/MAG_Snakemake_wf [Saheb Kashaf et al., 2021], MUFFIN [Van Damme et al., 2021], nf-core/mag [Krakau et al., 2021], and nbis-meta [, NBIS] have gained popularity with the promise of ease of use, reproducibility, and standardized protocols. These conveniences, however, often come at a cost of flexibility and transparency. Commonly developed and maintained by a single laboratory, these all-in-one workflows tend to have limited end-user customization potential. Easy-to-install workflows with well-chosen default settings can be suitable for researchers familiar with basic Unix commands, but can be analytic black boxes in nature, taking end-users from input to desired output with a one-line command. In this process, the intermediate results are implicitly obscured, preventing the end-user from integrating their domain expertise and allowing manual curation to play a role in data cleaning and analysis. In-depth intermediate results exploration, however, has been shown to be an integral part of ensuring MAG (metagenome-assembled genome) quality, especially in the reconstruction of circularized and (near-)complete microbial isolate genomes [Chen et al., 2019, Moss et al., 2020]. These factors combined left researchers with a world of metagenomic analysis pipelines generating substantially different results even in the same task with the same input [Sczyrba et al., 2017], a non-trivial obstacle for future metagenomic research.

Specifically, this workflow system should be highly flexible and transparent in terms of tool selection, parameter setting, and intermediate result exploration, while maintaining stability and a reasonable level of user-friendliness. In addition, this system should be comprehensive and scalable, able to cover most, if not all, aspects of metagenomic research needs, and with built-in Slurm integration for parallel processing on HPC systems, instead of focusing only on specific tasks such as MAG reconstruction. As metagenomics is an actively developing field with new tools being actively developed and potentially new analysis components being developed in the future, a future-facing workflow system should have the capacity to smoothly incorporate new tools and components without breaking the overall structure and also provide the environment for different tools to be benchmarked and compared.

With these objectives in mind, we developed CAMP, short for “Core Analysis Modular Pipeline”, a Snakemake-based fully modular workflow system designed for core metagenomic analyses with the above-mentioned features. A modular architecture, in contrast to the traditional monolithic design in most one-click pipelines, is one that designs workflows as an interconnected suite of modules, each responsible for one single analytic task (e.g. *de novo* assembly). While the concept of modular structure has been existing and there have been pipelines that claim to have adopted a “modular” architecture, such as nf-core/MAG [Krakau et al., 2021], FinnLab/MAG_Snakemake_wf [Saheb Kashaf et al., 2021], and MUFFIN [Van Damme et al., 2021], these pipelines are only modular insofar as having text. Most of these pipelines offer only limited end-user flexibility, having predefined workflows and tightly coupled configuration schema that need to be adhered to, such as in the case of nf-core/mag [Krakau et al., 2021]. Tweaking of these predefined structures often not possible without the modification of the internal code. In contrast, CAMP is fully modular with lightweight parameter files for each individual module, offering users the maximum freedom in designing their own pipelines. Each CAMP module is fully self-contained while sharing a common command-line interface and directory structure, balancing customization with usability.

Besides limited modularity, most of these abovementioned pipelines also suffer from a limitation of scope, focusing mainly on solving existing problems with focused goals such as assembly, binning, and profiling, and falls short in large-scale exploratory studies such that of MetaSUB [Danko et al., 2021]. CAMP, on the other hand, is designed for both current and future studies, both in its current comprehensiveness and in its flexible yet standardized design that allows unlimited number of future modules to be seamlessly incorporated. As an overview, we designed CAMP with the goal to establish a paradigm for next-generation metagenomic research by encapsulating the following features:

- **‘Set menu’-style computing to ‘a la carte’-style study design**: Switching from one-click pipelines to a modular analysis system allows the user to assemble the unique workflow for their specific study by downloading only the necessary modules.
- **Built-in soft pauses at the end of each module**: At the conclusion of each module, or ‘step’ in the workflow, the user can explore various semi-automated visualizations of their analytical results and apply their own knowledge base to enhance downstream analyses. This includes modifying downstream parameters from their default values.
- **Modules as benchmarking and comparison meta-tools:** A module can serve as a benchmarking ‘sandbox,’ akin to the Snakemake pipeline described by [Orjuela et al., 2022], where new methods can be seamlessly incorporated into a workflow and subsequently compared to other methods with similar objectives.
- **Compressed summary statistics tables**: The standardized output dataframes can serve as inputs for downstream machine learning ingestion.
- **Semi-guided visualizations**: Visualizations are essential for large-scale dataset analysis, hypothesis generation, and reliability assessments. To make effective judgment calls, it is important that understand what CAMP is doing at all intermediate steps.
- **Module template for future expansions**: Each module is based on a standard directory template. New modules for new analysis purposes can be easily set up from scratch using the `cookiecutter` command within a few hours. This process includes creating Conda environments and generating module-specific parameter and resource files.

## Methods

Each module is a GitHub repository containing core components organized in a standardized directory structure. This ensures consistency across the system, including the module directory (Box 1), working directory, parameterization, and input/output architectures. The core components include the `Snakefile, utils.py, ext/` directory, `parameter.yaml`, and `resource.yaml`, which are further customized for the module’s specific purpose. The design features of the modular system are outlined in Table 1, and available analysis modules are further described in Figure 1.

**Table 1.**
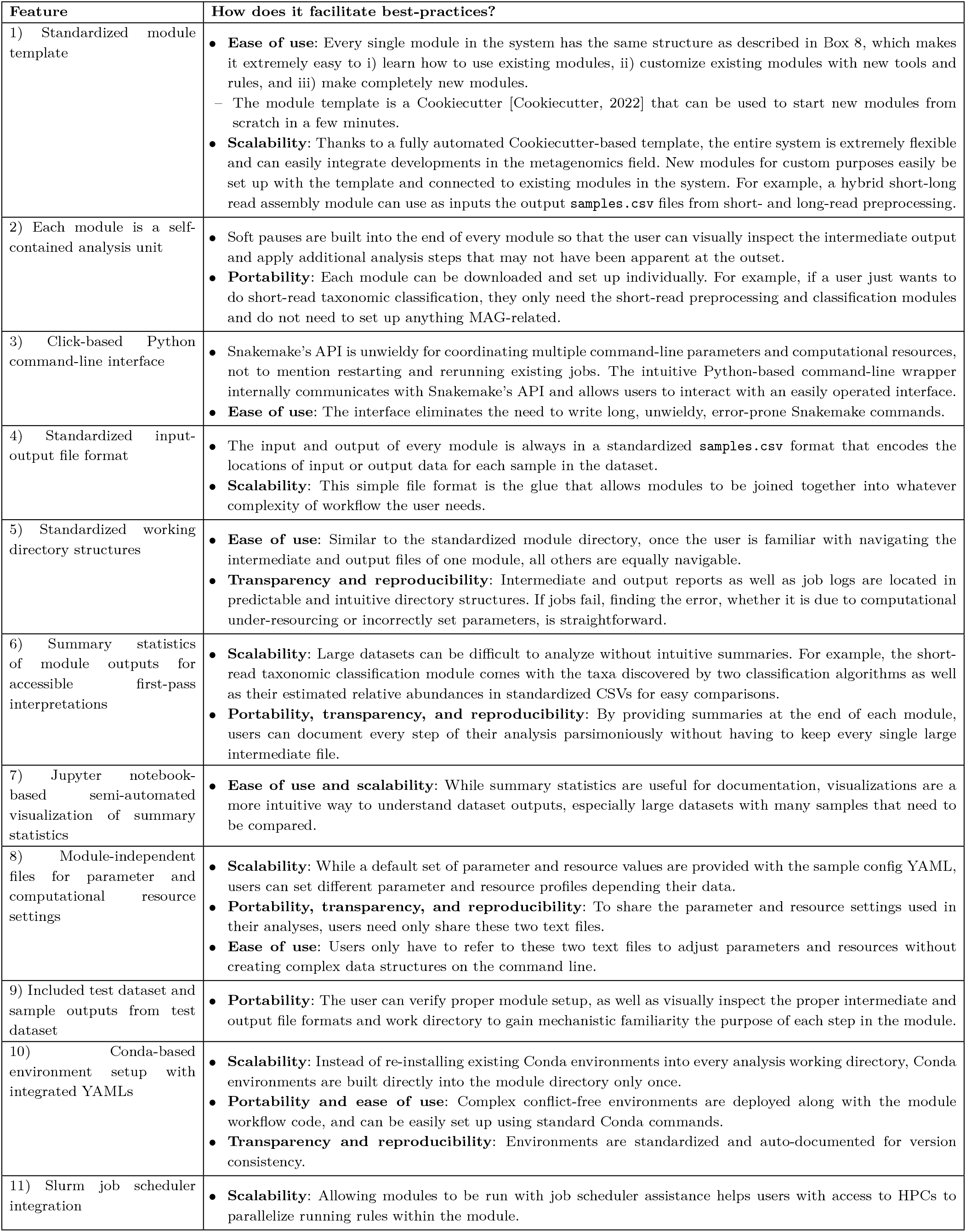
Each feature in the module system was designed to maximize scalability (in terms of dataset size as well as system distribution), portability (i.e.: compatibility with existing analysis environments), ease of use, and analysis transparency and reproducibility.

**Fig. 1.**
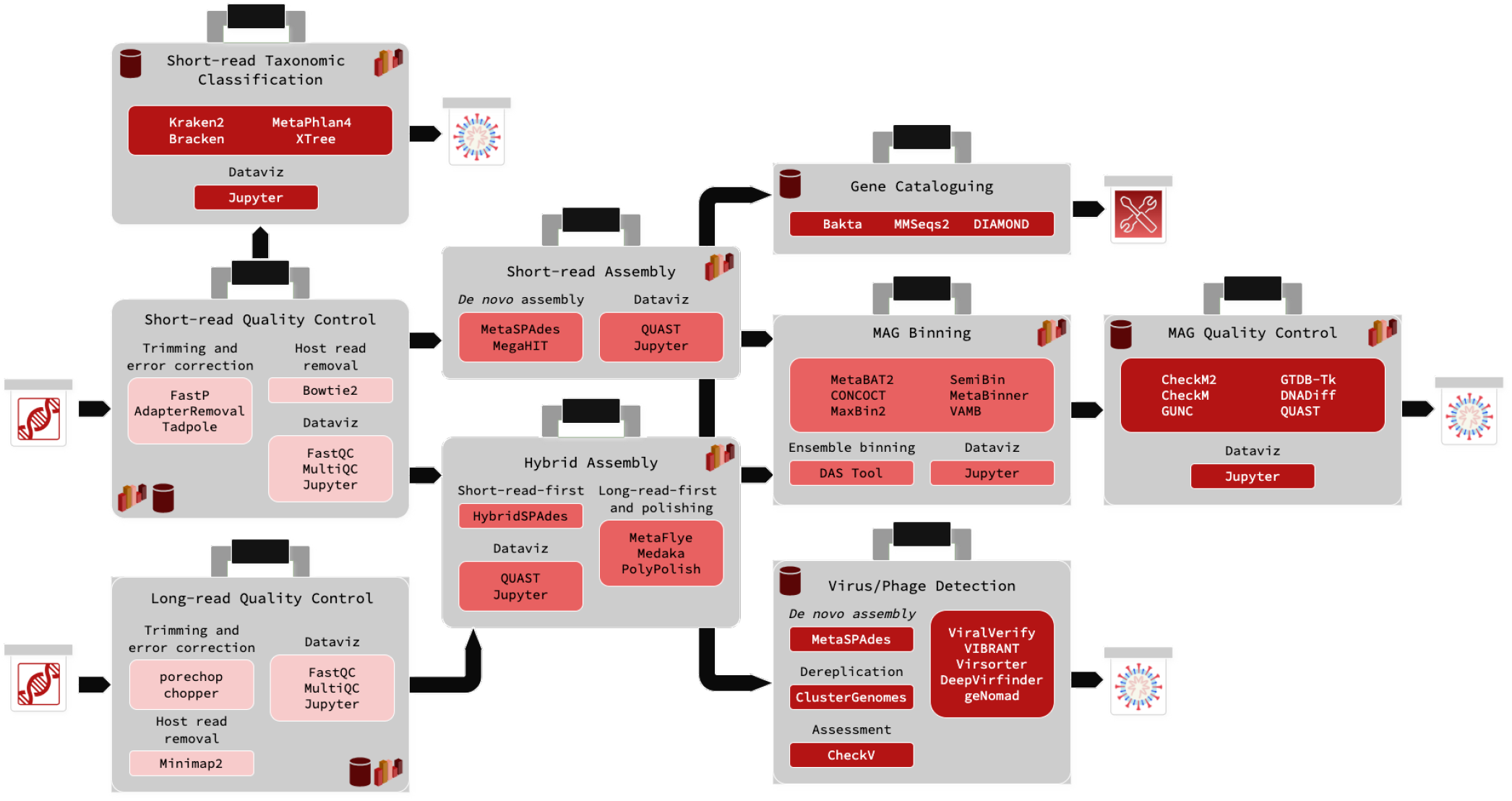
An overview of the available metagenomics analysis modules in the Core Analysis Modular Pipeline (CAMP). All modules share the same internal architecture, but wrap a different set of algorithms (shown to the left of each box) customized to its particular analysis goals. Modules that are typically the beginning of analysis projects are coloured light red, modules that are typically intermediate steps are coloured medium red, and modules that are typically terminal analysis steps are coloured dark red. Typical first inputs are indicated by the boxed DNA symbols, while terminal outputs that report some form of quality metric or taxonomic classification information are indicated by the boxed graphic bacteria. Functional profiling outputs are indicated with a boxed wrench and screwdriver. Modules that contain built-in dataviz notebooks contain a bar graph symbol, and modules that require database downloads contain a cylinder.

### Workflow Management System

While Nextflow [Di Tommaso et al., 2017] and Snakemake [Mölder et al., 2021] are both popular choices as workflow management systems for computational biology analysis, we opted for Snakemake due to its general readability and accessibility, particularly for researchers accustomed to using Python. Snakemake’s syntax is minimal and resembles standard Python, making workflows easy to understand, develop, expand, and troubleshoot. Although working with Snakemake’s wildcards may pose a learning curve, its directives (e.g., output, log) and ability to specify file names explicitly generally result in output directories that are easy to navigate and conducive to workflow reruns. Nextflow also offers many strengths as a workflow management system (WMS), with better support for cloud deployment natively as well as nf-core, a repository of curated analysis pipelines developed by the community at large [Ewels et al., 2020].

### Environment Management

We have opted not to implement the modular system with Docker [Merkel, 2014] or Singularity [Kurtzer, 2018], because many users of high-performance compute clusters (HPCs), lack root access privileges and thus cannot utilize containerized workflows. Instead, we have included conflict-free recipes in each module directory, each of which sets up a Conda environment directly. This allows the same environment to be used for all datasets processed using the module [conda contributors]. To ensure reproducibility and interoperability within the module environment, each Conda recipe has hard-coded algorithm versions and is designed specifically for Unix-based operating systems. In response to Anaconda’s licensing policy as of March 2024, the `defaults` channel have been phased out of each module’s Conda recipe. This allows users to recreate environments from YAML files without automatically pulling packages from the proprietary channel, facilitating a smoother transition for those shifting to community-supported channels like `conda-forge` that remain unrestricted.

### Modular System Use-cases

The primary strengths of a modular system are its flexibility and extensibility, complemented by the interdependency of the tools. This architecture enhances accessibility and simplifies maintenance, enabling a longer lifespan. Such a design is accessible not only to the developers of the module template and core modules but also to anyone with fundamental command-line, Python, and Conda development skills. Below are a few use cases:

1. A user possessing a short-read dataset and seeking to conduct an initial analysis with short-read taxonomic classification can download two modules: “short-read preprocessing” for the removal of low-quality and erroneous bases, and “short-read taxonomic classification” for the generation of a unified report consolidating discovered taxa from two classification algorithms. Simplifying the learning process, the user interfaces with a single command-line interface (CLI), which remains consistent across all modules.
2. The user, intending to expand their analysis to include gene annotation, can effortlessly do so by downloading and executing two additional modules: “short-read assembly” and “gene cataloguing.” No additional onboarding time is necessary in terms of interface learning. Moreover, the input for short-read assembly remains consistent with that of short-read taxonomic classification, utilizing the `samples.csv` file containing paths to preprocessed short reads.
3. The user, aiming to broaden their analysis to include MAG reconstruction, can seamlessly achieve this by downloading the binning and MAG quality-checking modules. Utilizing the output of short-read assembly, specifically the samples.csv file containing paths to the *de novo* assemblies, as their input, no additional onboarding time is required to learn new interfaces.
4. 4.A user who has developed a new short-read taxonomic classification tool can leverage the existing short-read taxonomic module as a sandbox environment. They can write a custom Snakemake rule to execute their tool, implement a function to harmonize the output format with the existing merged report, and conduct benchmarking analyses to assess the strengths and weaknesses of their tool.
5. 5.A user aiming to study extra-chromosomal elements, given the absence of an existing module for this purpose, can create a new blank module utilizing the Cookiecutter template. They can then populate it with the requisite Snakemake rules and customize the remaining demo configurations to suit their specific objectives. Leveraging the standardized input-output format ensures seamless integration of this new extra-chromosomal elements module with the broader module system, facilitating reuse by both the developer and the wider research community.

### Currently Available Modules

There are ten modules available, and seven are used in this proof-of-concept analysis to accomplish one of three purposes: i) short-read taxonomic classification, ii) MAG reconstruction and quality-checking, and iii) virus and phage inference from short-read sequencing datasets (Figure 1). There are several modules currently under active development, including Oxford Nanopore long-read preprocessing, mOTUs, and decontamination. The algorithms included for each module were chosen based on previously documented performance in benchmarking studies, as well as code stability and overall workflow compatibility.

### General-Purpose Analysis

#### Module 1: Short-read Preprocessing

Raw sequencing datasets are filtered for low-quality bases, low-complexity regions in reads, extremely short reads, and adapter content using fastp [Chen et al., 2018]. Reads can optionally be deduplicated. If host read removal is selected, trimmed and filtered reads are mapped using Bowtie2 and Samtools with the ‘very-sensitive’ flag to the host reference genome (here, the human reference genome assembly GRCh38), and mapped reads removed [Langmead and Salzberg, 2012, Li et al., 2009]. As a last-pass, BayesHammer or Tadpole is used to correct sequencing errors [Nikolenko et al., 2013, Bushnell, 2014]. FastQC and MultiQC are used to generate overviews (e.g. parameters such as per-base quality scores, sequence duplication levels) of processed dataset quality [Andrews, 2010, Ewels et al., 2016].

#### Module 2: Nanopore Long-Read Quality Control

Raw sequencing datasets are trimmed using PoreChop [Wick, 2018], and then low-quality bases are filtered out using chopper [Coster and Rademakers, 2023]. Host reads are optionally removed using Minimap2 [Li, 2018]. FastQC and MultiQC are used to generate overviews (e.g. parameters such as per-base quality scores, sequence duplication levels) of processed dataset quality [Andrews, 2010, Ewels et al., 2016].

### *De novo* Assembly

#### Module 3: Short-read Assembly

The processed sequencing reads can be assembled using MetaSPAdes (with optional flags for metaviral and/or plasmid assembly also available), MegaHIT, or both [Nurk et al., 2017, Li et al., 2015]. For the purposes of this study, only MetaSPAdes was used. The assembly is subsequently summarized using QUAST [Gurevich et al., 2013].

#### Module 4: Hybrid Assembly

For short-read-first hybrid assembly, processed short sequencing reads are assembled with hybridmetaSPAdes, with the draft assembly polished by processed long reads [Antipov et al., 2016]. For long-read-first hybrid assembly, processed long reads are assembled with MetaFlye [Kolmogorov et al., 2020], before being corrected first with the long reads using Medaka [Ltd., 2019] and then with short reads using PolyPolish [Bouras et al., 2024]. The assemblies are subsequently summarized and compared using QUAST [Gurevich et al., 2013].

### MAG Inference and Quality-Checking

#### Module 5: MAG Binning

Processed sequencing reads are mapped back to the de novo assembled contigs using Bowtie2 and Samtools. This read coverage information, along with the contig sequences themselves, are used as input for the following binning methods: MetaBAT2, CONCOCT, SemiBin, MaxBin2, VAMB, and MetaBinner [Kang et al., 2019, Alneberg et al., 2014, Pan et al., 2022, Nissen et al., 2021, Wu et al., 2016, Wang et al., 2023]. The sets of MAGs inferred by each algorithm are used as input for DAS Tool, an ensemble binning methods, to generate a set of consensus MAGs scored based on the presence/absence of single-copy genes (SCGs) [Sieber et al., 2017]. Some of the contig pre-processing scripts were adapted from the MAG Snakemake workflow [Saheb Kashaf et al., 2021].

#### Module 6: MAG Quality-Checking

The consensus refined MAGs are quality-checked using an array of parameters. CheckM2 calculates completeness and contamination based on the annotated gene content of a MAG [Chklovski et al., 2023]. CheckM1 calculates per-MAG short-read coverage and strain heterogeneity based on the proportion of presence of multiple copies of single-copy marker genes that pass a high amino acid identity threshold [Parks et al., 2015]. gunc is also used to assess contamination [Orakov et al., 2021]. MAGs are classified using GTDB-Tk, which relies on approximately calculating average nucleotide identity (ANI) to a database of reference genomes [Chaumeil et al., 2019]. For MAGs with a species classification, their contig content is compared to the species’ reference genome and genome-based completion, misassembly, and non-alignment statistics calculated using dnadiff and QUAST [Marçais et al., 2018, Gurevich et al., 2013]. Each MAG’s gene content, with an emphasis on tRNA and rRNA genes in accordance with MIMAG genome reporting standards [Bowers et al., 2017], is summarized using prokka [Seemann, 2014].

The overall score per MAG was calculated using CheckM2-calculated completeness and contamination, and the contiguity metric N50: completeness *−* 5 *×* contamination + 0.5 *×* log(*N* 50). This equation was adapted from dRep’s overall score equation, which is based on completeness, contamination, contiguity, strain heterogeneity, and genome size, and [Olm et al., 2017]. dRep used this score to select a representative genome from a cluster of genomes with similar sequences. Various versions of this equation exist, with most variants setting the coefficients for strain heterogeneity and genome size to 0 [Williams et al., 2024, Gurbich et al., 2023, Parks et al., 2017].

### Other Analysis Goals

#### Module 7: Short-read Taxonomic Classification

The processed sequencing reads can be classified using MetaPhlan4, Kraken2/Bracken, and/or XTree [Wood et al., 2019, Lu et al., 2017, Blanco-Miguez et al., 2022, GabeAl, 2022]. XTree, formerly referred to as UTree, is additionally included as an experimental short-read classification option, but this study focuses on comparing the taxa discovered by the two published field-standard tools-Kraken2 and MetaPhlan4. To estimate the relative abundance of a taxon, MetaPhlan4 calculates marker gene coverage and Bracken calculates the proportion of reads assigned to a taxon with k-mer uniqueness-based scaling [Blanco-Miguez et al., 2022]. Since each of these output reports are of different formats, the raw reports from each algorithm are standardized in format for easier comparisons downstream.

#### Module 8: Virus/Phage Inference

The processed sequencing reads are assembled with MetaSPAdes, and viral contigs are subsequently identified using the output assembly graph and ViralVerify [Nurk et al., 2017, Antipov et al., 2020]. Contigs containing putative viral genetic material are also identified using VIBRANT, VirSorter2, DeepVirFinder, and geNomad [Kieft et al., 2020, Guo et al., 2021, Ren et al., 2020, Camargo et al., 2024]. The aggregated lists of contigs from the three inference methods is dereplicated using VirClust [Moraru, 2023] and merged with the ViralVerify list, and the overall quality of the putative viruses is assessed using CheckV [Nayfach et al., 2021].

#### Module 9: Gene Cataloguing

Open reading frames (ORFs) are identified in the de novo assembly using Bakta, and clustered using MMSeqs [Schwengers et al., 2021, Hauser et al., 2016]. Genes are identified from these ORFs by alignment to the DIAMOND database to obtain the functional profile of the sample [Buchfink et al., 2014].

#### Module 10: Decontamination

The decontamination module is still under construction. A feature table of relative abundances (e.g. operational taxonomic units (OTUs), taxa, metagenome-assembled genomes) is provided to Decontam and Recentrifuge, each of which estimates contamination from feature abundances either within or between samples respectively [Davis et al., 2018, Martí, 2019].

#### Module 11: mOTUs Profiling

The long-read profiling module is still under construction. It wraps mOTUs [Ruscheweyh et al., 2021], which estimates the relative abundance of taxa from a short- or long-read sequencing dataset, and calls single-nucleotide variants from marker genes.

### Comparison with Other Workflows

To further delineate what sets CAMP apart, we compare CAMP with five other pipelines commonly used in metagenomics analysis: ATLAS Kieser et al. [2019], FinnLab/MAG_Snakemake_wf [Saheb Kashaf et al., 2021], MUFFIN [Van Damme et al., 2021], nf-core/mag [Krakau et al., 2021], and nbis-meta [, NBIS] (Table 2). It is worth pointing out that these pipelines, though used in an array of modern metagenomic analysis procedures, are mostly MAG-focused, as many of the names suggest. CAMP offers a distinctly different philosophy of project design, which envisions a comprehensive and extensible system capable of supporting all core and extant analyses in current and future metagenomic research.

**Table 2.** A comparison of features found across several recently published metagenomics analysis pipelines and CAMP. The symbol ‘+’ indicates that the feature has been implemented in the pipeline, with more ‘+’ indicating extensiveness. Conversely, the ‘-’ indicates that the feature is absent. The ‘MAG Binning’ row is formatted as X+Y, where X refers to the number of MAG binning algorithms and Y indicates the presence of a consensus binning tool.

| Feature | CAMP | ATLAS | MAG Snakemake wf | MUFFIN | nf-core/mag | nbis-meta |
| --- | --- | --- | --- | --- | --- | --- |
| Language | Snakemake | Snakemake | Snakemake | Nextflow | Nextflow | Snakemake |
| Modular design? | + | - | - | - | - | - |
| Install testing? | + | + | - | + | + | - |
| Containerized? | - | + | + | + | + | + |
| Documentation? | ++ | + | ++ | ++ | ++++ | + |
| Data Visualization | + | - | + | - | - | - |
| Nanopore Long-Read QC | + | - | - | + | + | - |
| Hybrid Assembly | + | - | - | + | + | - |
| Co-assembly and co-binning | - | - | + | - | + | - |
| MAG Binning | 6+1 | 2+1 | 3+1 | 3+1 | 3+1 | 3 |
| Short-read Taxonomic Classification | + | - | - | - | + | + |
| Virus/Phage Detection | + | - | - | - | + | - |

For example, aside from CAMP, only nf-core/mag offers both short-read taxonomic classification and virus/phage detection-two additional analysis results unrelated to MAG binning and quality-checking [Krakau et al., 2021]. Even with the respect to core MAG-related tasks, however, we have demonstrated our commitment to building modules that are easily extensible. CAMP incorporates 6 different binning algorithms, which is far and away the maximum of 3 incorporated by other workflows (Table 2). In addition to the classic options such as MetaBAT2 [Kang et al., 2019], CONCOCT [Alneberg et al., 2014] and MaxBin2 [Wu et al., 2016] which are commonly used in other pipelines, the binning module includes newer machine learning-based binning tools such as VAMB [Nissen et al., 2021] and SemiBin [Pan et al., 2022]. In the Results section below, we further demonstrate why diversity on perspective in MAG binning is necessary. Different binning strategies return different numbers of MAG bins, many of which are of questionable quality that would not have been identified using the classic options alone. For large-scale studies that want to balance MAG quality with speed, CAMP goes above and beyond to provide the most robust analysis results.

## Results

### Proof-of-Concept and Data

To demonstrate CAMP’s applicability for large-scale metagenomics analysis and inter-sample comparison, we have applied seven modules to a randomly selected set of 10 samples from the International Metagenomics and Metadesign of Subways and Urban Biomes (MetaSUB) dataset, which consisted (at time of [Danko et al., 2021]’s publication) of 4,728 samples of microbiomes collected from urban transit system surfaces between 2015 - 2017. Sample collection, metagenomic DNA extraction, and sequencing information can be found in [Danko et al., 2021] and the GeoSeeq repository. The raw sequencing sequencing data can be found under the SRA accession ID PRJNA732392, as well as the GeoSeeq repository. The 10 randomly chosen samples were collected from subway train and transit station surfaces from three of the four boroughs of New York that are connected by subway transit lines-Manhattan, Queens, and Brooklyn (no samples from the Bronx were in the subset, Table 3). Most of the samples were obtained from transit station surfaces in the winter of 2014, and generally contain between 2 - 6 million paired-end reads.

**Table 3.**
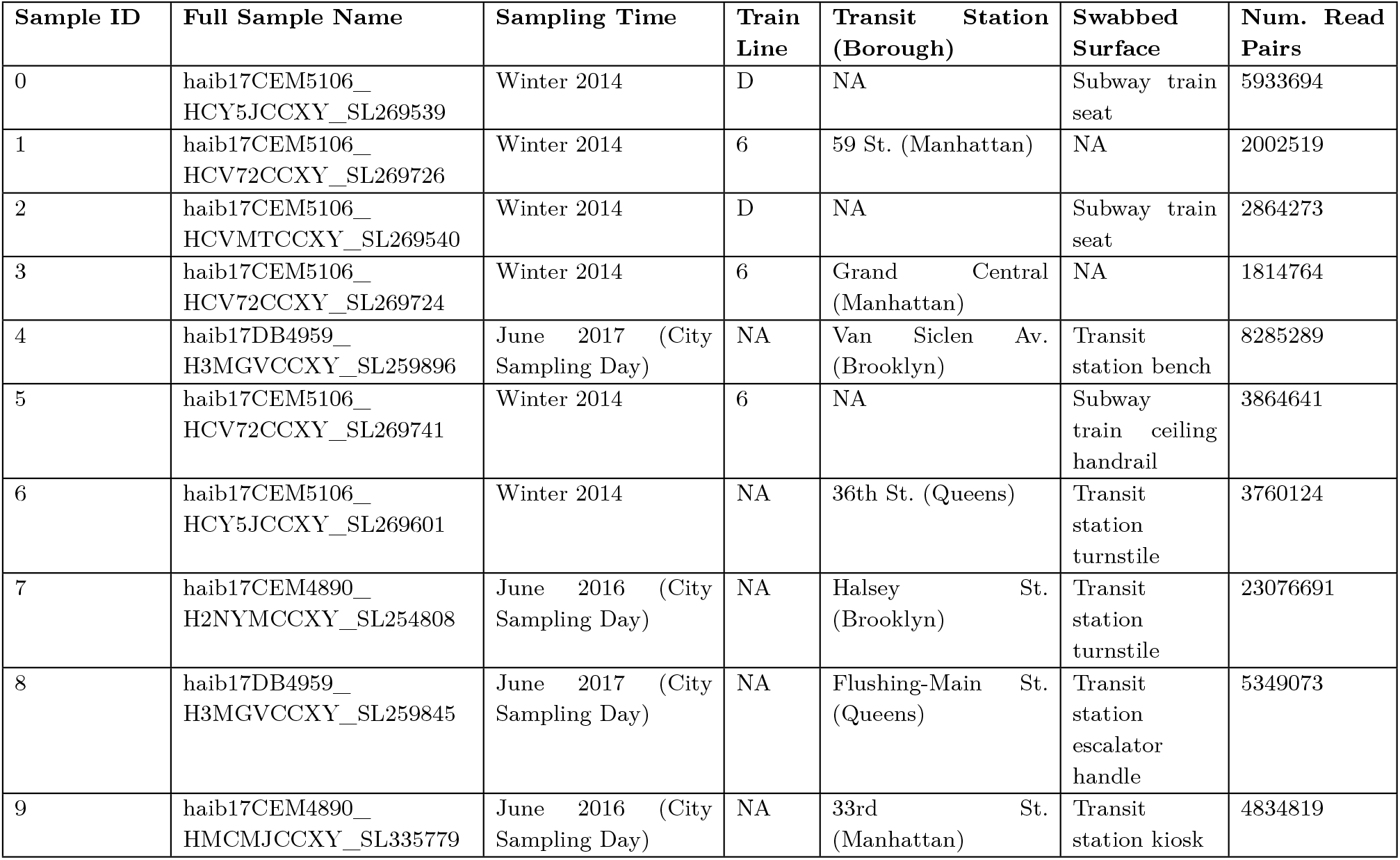
Ten randomly chosen urban microbiome samples from the MetaSUB New York dataset along with metadata such as their dates and locations of origin.

### Short-Read Quality Control

Between 66.6 - 93.8% of short sequencing reads and 65.3 - 89.8% of sequencing bases were retained after low-quality base-trimming, adapter trimming, host (human genome GRCh version 38) genome removal, and error correction steps were applied to each of the 10 samples (Figure 2, Supp. Table 1). The largest reduction in dataset size occurred at the `fastp` low-quality read removal step, with host read removal by Tadpole also eliminating some sequence content (Figure 2A). No reads and bases were lost at the adapter step despite adapters being present in the pre-flight MultiQC check, indicating that all of the adapter bases were trimmed by `fastp`.

**Fig. 2.**
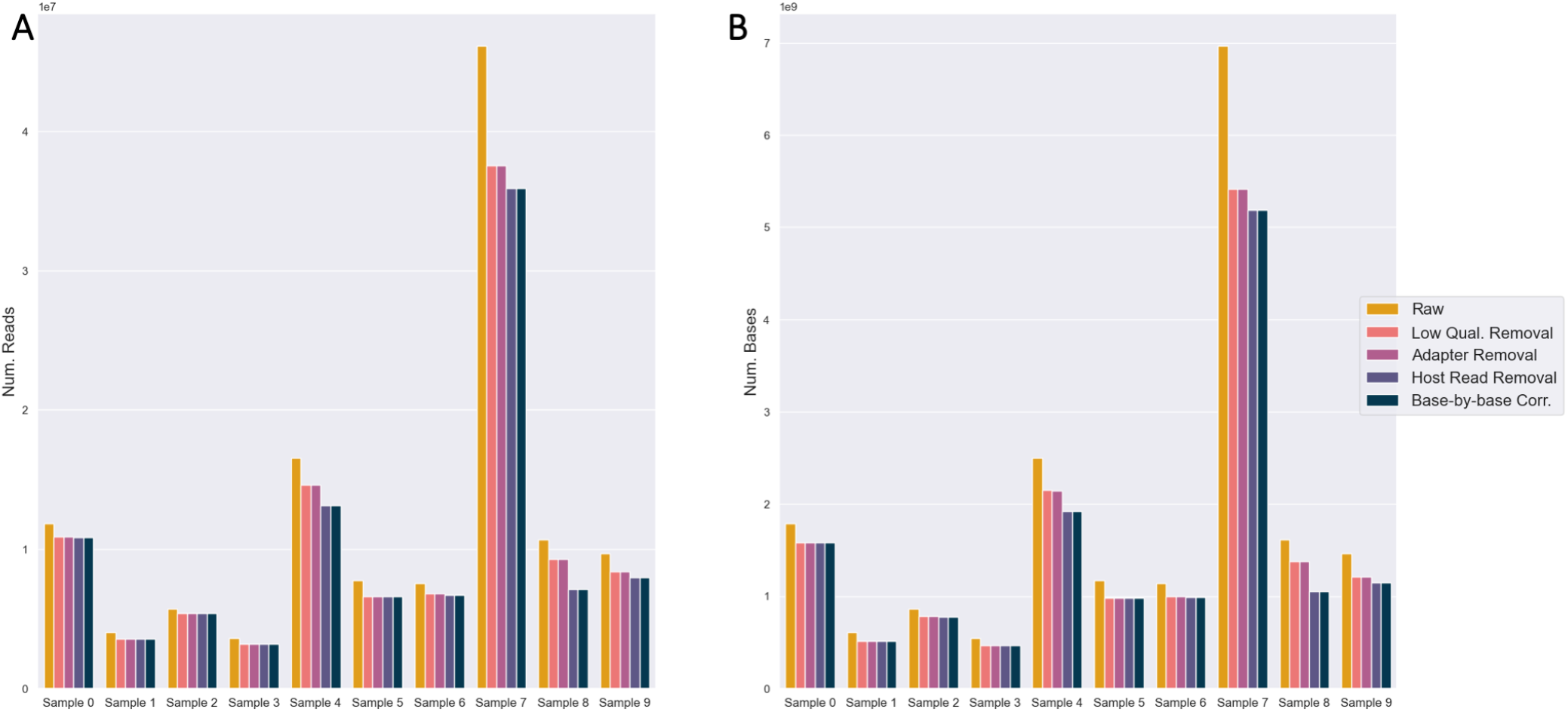
Quality control and preprocessing of short-read sequencing data. Counts of (A) reads and (B) bases retained after each preprocessing step across all samples. The steps include low-quality base trimming, adapter removal, host genome removal, and error correction.

After preprocessing, most samples passed most MultiQC quality metrics. For the reverse reads of samples 8 - 9 and samples 5 - 7, there were outstanding warnings for the per-tile sequencing quality (example in Supp. Figure 1A) and per-base sequence content (example in Supp. Figure 1B). Upon checking the FastQC plots for samples 8 - 9, in both cases, the tile failed at one position in particular (towards the end), not across all bases (Supp. Figure 1A). Since the average per-base quality is generally high, no further trimming was pursued. For samples 5 - 7, the ACGT content was uneven for the first 10 bases (Supp. Figure 1B). It has been demonstrated previously in RNASeq datasets that 5’ nucleotide biases can be caused by steps in library preparation [Hansen et al., 2010]. Since all adapters were removed and overall sequencing quality was high, no further trimming was pursued.

### Short-Read Assembly

The distribution of assembly sizes mostly follows the distribution of short-read dataset size, with the exception of Sample 4 (Figure 3A, Figure 2B). Although the size of Sample 7 is nearly 3X larger than sample 4 (in terms of both numbers of reads and bases), Sample 4 has the largest assembly by far in terms of both numbers of contigs and bases (Figure 3). MultiQC reported that Samples 4 and 7 contained 7.4% and 10.9% duplicated reads overall, which accounts minimally for the difference. Sample 4 may contain a much lower proportion of PCR duplicates. As shown later by both short-read taxonomic classification and MAG binning, sample 4 does not contain appreciably fewer species than sample 7. The largest, average, and median contig sizes are similar, though there is sizable variance in terms of assembly contiguity as measured by N50 (Supp. Table 2).

**Fig. 3.**
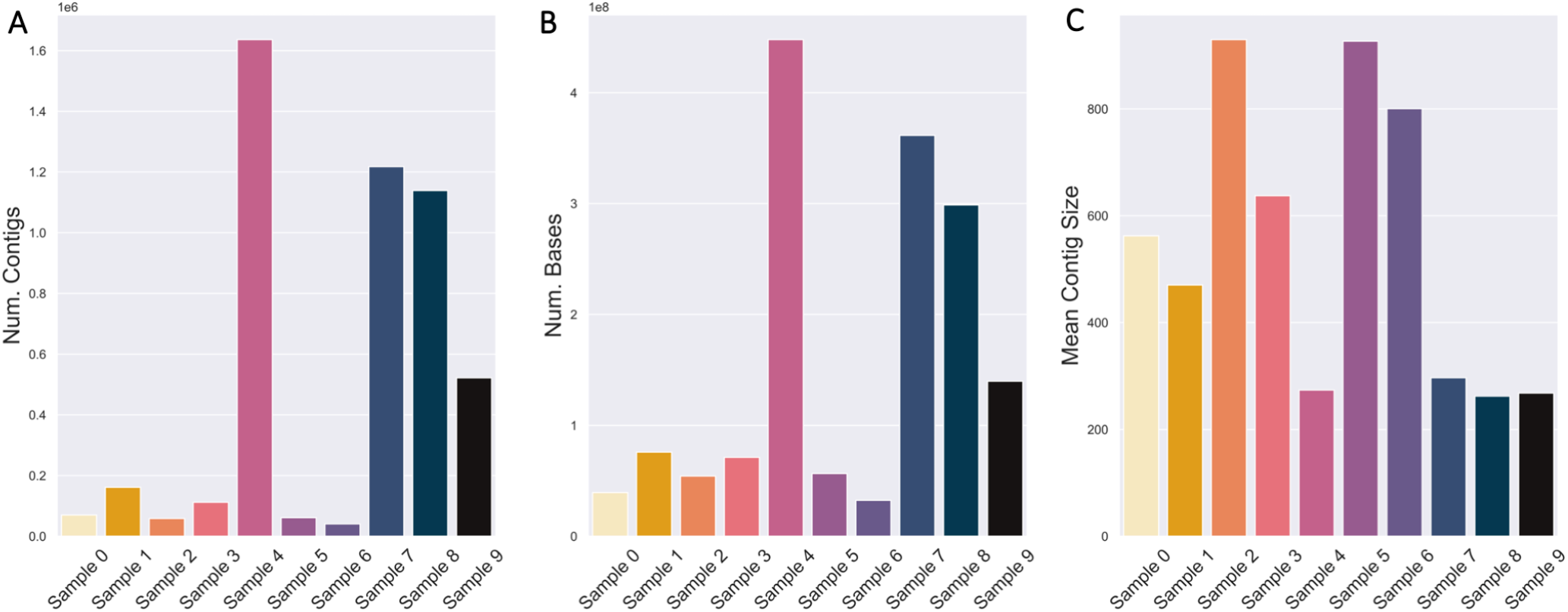
*De novo* assembly sizes generally correlate with short-read sequencing dataset sizes. (A) The number of contigs and (B) number of sequencing bases in each sample’s assembly, as well as the (C) mean contig size.

The vast majority of contigs in all *de novo* assemblies are extremely small and fall below the size cutoffs of most MAG binning algorithms (two examples in Table 4). Smaller contigs are more likely to contain low-quality bases and have irregular coverage [Hofmeyr et al., 2020], and are also more likely to originate from low-abundance organisms [Vicedomini et al., 2021]. To retain only high-quality sequencing information for binning, we selected a MAG binning minimum contig cutoff of 2500 base-pairs. Only a small fraction of the assemblies are of contigs larger than 50 Kbp (0.2% at maximum), which is approximately 1% of the size of the average bacterial genome [diCenzo and Finan, 2017].

**Table 4.** With a minimum contig size threshold of 2500 bp, a tiny minority of contigs- between 0.1-0.5%-will be kept for MAG binning.

| Sample | Min. Contig Size | Num. Contigs (Prop.) | Num. Bases (Prop.) |
| --- | --- | --- | --- |
| 4 | 0 bp | 1636542 (1.000) | 447963811 (1.000) |
| 4 | 2,500 bp | 1872 (0.001) | 10832446 (0.024) |
| 4 | 50,000 bp | 12 (0.000) | 1000212 (0.002) |
| 4 | 100,000 bp | 3 (0.000) | 342030 (0.001) |
| 7 | 0 bp | 1217385 (1.000) | 361500456 (1.000) |
| 7 | 2,500 bp | 6605 (0.005) | 53796944 (0.149) |
| 7 | 50,000 bp | 94 (0.000) | 11802085 (0.033) |
| 7 | 100,000 bp | 41 (0.000) | 8342312 (0.023) |

### MAG Binning

Aside from VAMB and samples 7 - 9, the number of MAGs inferred by each binner (including the MAG inference aggregator and ensemble binner DAS Tool) was generally within a factor of two of each other (Figure 4A). In general, the more MAGs inferred by a binner, the smaller the inferred MAGs on average in terms of the numbers of sequencing bases (Figure 4A,C). The three binners that do not use single-copy marker genes to estimate the number of MAGs-MetaBAT2, CONCOCT, and SemiBin-tend to infer a larger number of MAGs (Figure 4A) [Kang et al., 2019, Alneberg et al., 2014, Pan et al., 2022, Nissen et al., 2021, Wu et al., 2016, Wang et al., 2023, Sieber et al., 2017]. SemiBin only uses single-copy marker genes to further partition MAGs with an average of *>* 1 copy per gene [Pan et al., 2022]. While MaxBin2 and MetaBinner generally inferred the same number of MAGs, MaxBin2 MAGs generally contain many more contigs but are similar in overall size (as counted by base pairs). This can be explained by MetaBinner’s stringent post-clustering contig reassignment step; MetaBinner uses a MetaWRAP-like consensus method to remove contigs that are not robustly associated with cluster centroids [Wang et al., 2023].

Despite having the largest assembly, only 4 MAGs were inferred from sample 4 by DAS Tool (Table 5). Furthermore, the size of the MAGs is approximately 2 Mbp, which is smaller than the average bacterial genome (Figure 4C) [diCenzo and Finan, 2017]. The largest number of MAGs was inferred from sample 2 (Table 5), which was the third smallest dataset and assembly (by the number of bases, Figure 2B, Figure 3B). However, sample 2 also had the largest average contig size, so more contigs were included in the binning process after applying a minimum size threshold of 2500 bp (Figure 3C).

**Table 5.** The number of MAGs inferred by the 6 binning algorithms and ensemble-aggregated by DAS Tool, as well as the number of MAGs that can be classified by GTDB-Tk.

| Sample | Num. DAS Tool MAGs | Num. Classifiable MAGs | Num. Species-Classifiable MAGs |
| --- | --- | --- | --- |
| 0 | 4 | 4 | 3 (75%) |
| 1 | 4 | 4 | 4 (100%) |
| 2 | 8 | 8 | 4 (50%) |
| 3 | 6 | 6 | 5 (83.3%) |
| 4 | 4 | 4 | 3 (75%) |
| 5 | 7 | 7 | 5 (71.4%) |
| 6 | 5 | 5 | 5 (100%) |
| 7 | 3 | 3 | 1 (33.3%) |
| 8 | 2 | 2 | 2 (100%) |
| 9 | 1 | 1 | 1 (100%) |

Aside from VAMB, DAS Tool generally inferred the fewest MAGs, though they are usually the largest and similar to the average size of a bacterial genome (Figure 4C) [diCenzo and Finan, 2017]. In terms of overall assembly utilization, DAS Tool MAGs contained between 91.3 - 99.7% of usable bases in contigs >2500 bp.

### MAG QC

Most of the inferred MAGs (27/44, 61.3%) across all samples were high-quality, according to a slightly modified version of the MIMAG standards naming conventions (Figure 5R) [Bowers et al., 2017]. MAGs that are high-quality but missing the requisite rRNA and tRNA genes are still called high-quality, and those with are called ‘draft’ instead. There were no MAGs that contained all of the rRNA and tRNA genes required for ‘draft’ status.

Similarly, most of the MAGs (34/44) had CSSes that fell under the GUNC threshold for allowable taxonomic chimerism. Of the low-quality 5/44 (9.2%) MAGs, 2/5 had low CheckM2-estimated contamination and gunc-estimated clade separation score (CSS) and were still classifiable to the species level. The remaining low-quality high contamination MAG (CheckM2: 10.81, gunc: 0.97) was classified to the genus Pseudomonas. Given the high CSS, the MAG may be a chimeric bin of contigs from multiple genera in the *Pseudomonadaceae* family. CheckM2-estimated completeness was *≤* 50% for only a single MAG, and contamination was *≥* 10 for three (Figure 5B). Of the species-classifiable MAGs, contamination was at maximum 5, but some had high CSS, up to 1.

Recall that only a small fraction of contigs were larger than 50 Kbp (Table 4). The average size of the contigs in the MAGs of samples 4 and 7 are 9.7 and 18.3 Kbp respectively (Figure 4C). The median NA50 of the classifiable MAGs are 8.5 and 35.3 Kbp respectively (Figure 5M), indicating that most of the assemble-able and mappable content of a MAG is concentrated within a small number of large contigs. The rest of the associated reference genome may be difficult to assemble and may have different sequence properties than the rest of the contigs. The use of BLAST or a taxonomic classifier (e.g. Kraken2, MetaPhlan4, CAT [Wood et al., 2019, Blanco-Miguez et al., 2022, Cambuy et al., 2016]) to classify short contigs may be more appropriate, as they may be small fragments from low-abundant species too poorly covered by sequence content to bin into MAGs.

**Fig. 4.**
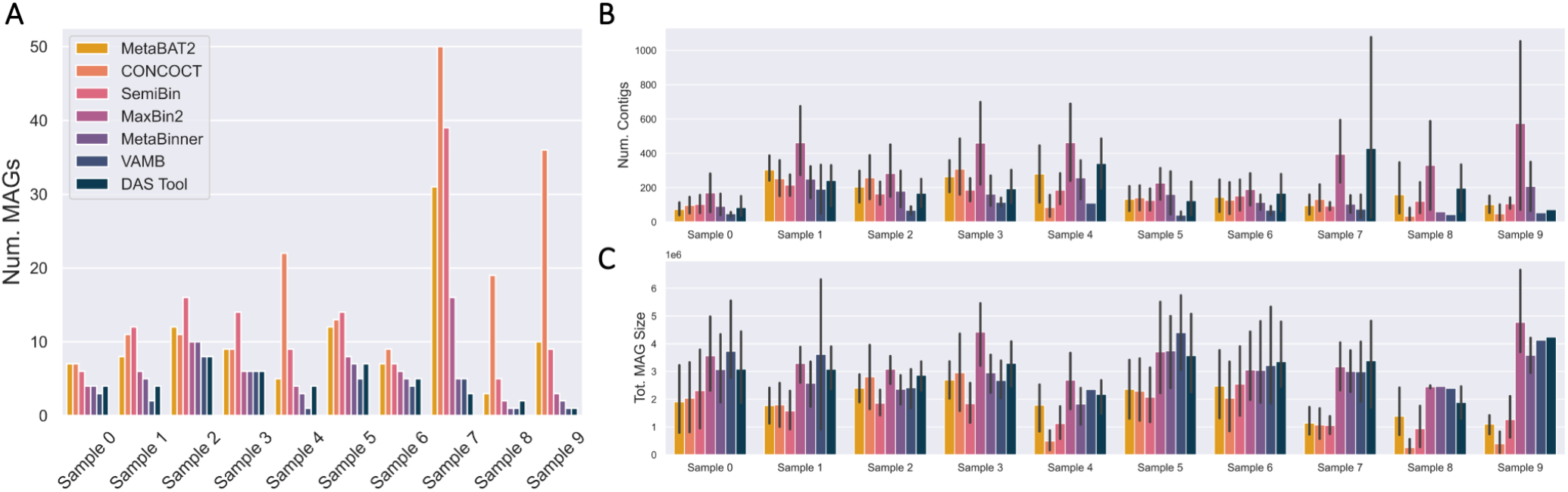
Most MAG binning algorithms infer a consistent number of MAGs, with the exceptions of samples 4, and 7 - 9 and VAMB. (A) Number of MAGs inferred by each binning algorithm across samples. (B) The numbers of contigs per (C) Total MAG size inferred for each sample and comparison across binners (bars indicate standard deviation).

**Fig. 5.**
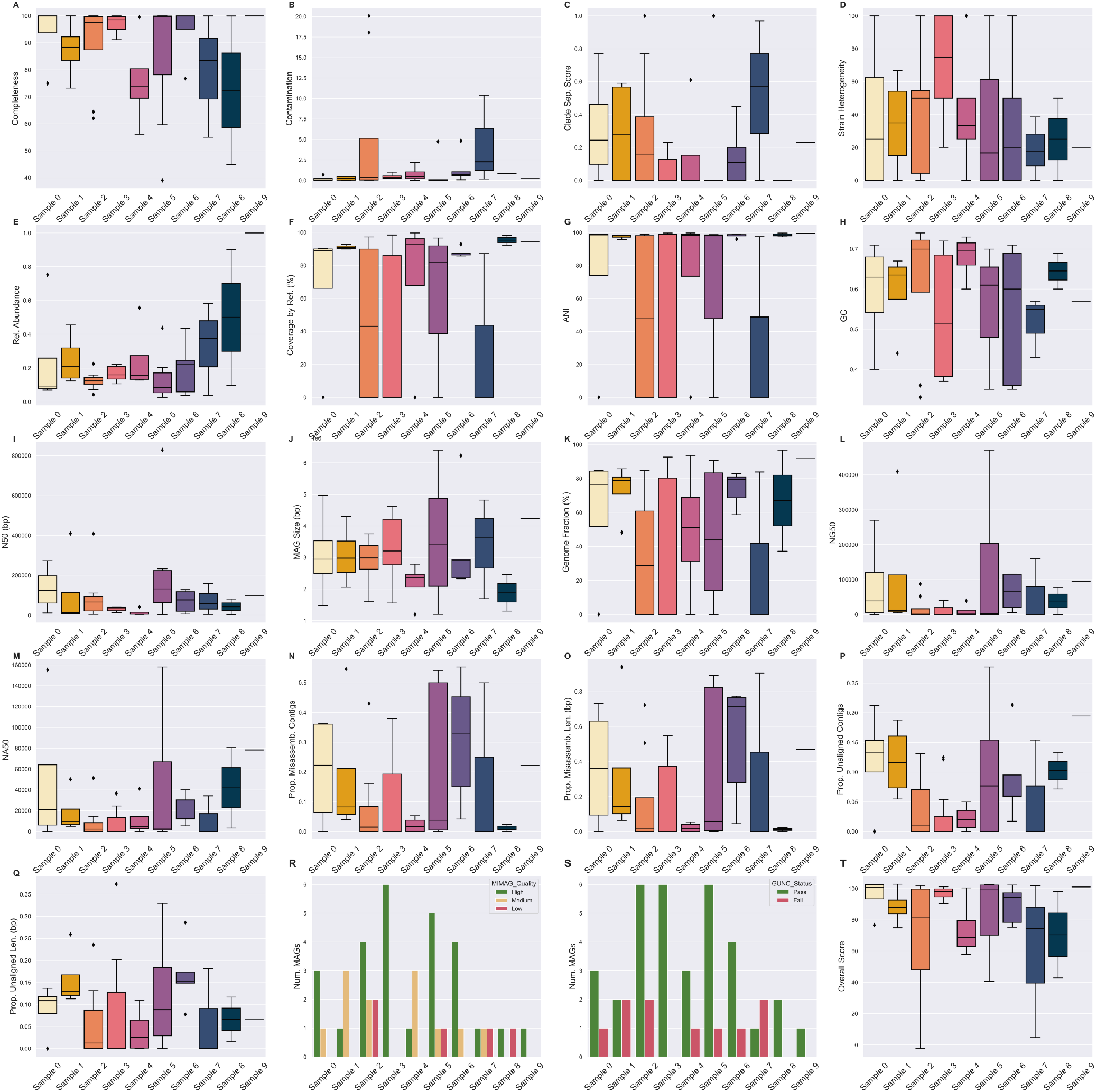
Quality assessment metrics of DAS Tool-inferred MAGs across all samples. The median of plot’s metric across all of the MAGs inferred from that sample is indicated with a line across the box. (A) Completeness, (B) contamination, (H) GC, (I) N50, and (J) MAG size were reported by CheckM2. (C) Clade separation score was reported by gunc. (D) Strain heterogeneity and (E) relative abundance were reported by and calculated from CheckM1 respectively. (F) MAG coverage by the GTDB-Tk classified reference genome and (G) average nucleotide identity (ANI) to that genome were reported by dnadiff. Metrics (K - Q) were reported by QUAST comparing each MAG to the GTDB-Tk classified reference genome. (T) was calculated from the MAG’s completeness, contamination, and N50.

The median relative abundance of each MAG per sample is generally between 10 - 20%, with half of the distributions having some right skew i.e.: there are a few MAGs where the sample relative abundance is much higher than the average (Figure 5E), which generally agrees with the expected log-normal distribution of bacterial relative abundances in a sample [Hill et al., 2003, Prost et al., 2021].

Completeness is highly correlated with QUAST-estimated genome fraction (the fraction of the reference genome that the MAG aligned to) (Figure 6A). However, contamination and CSS do not correlate with the proportion of unaligned sequence material (Figure 6B, C). Though both CheckM2 and gunc use a MAG’s protein content to estimate contamination, they differ in terms of application and thus interpretation. CheckM2 uses a neural network trained from almost protein-annotated 5,000 RefSeq bacterial genomes [Chklovski et al., 2023]. gunc classifies the candidate proteins and measures the taxonomic entropy within a contig and the MAG overall [Orakov et al., 2021]. If a MAG is high in CheckM2-estimated contamination but low in gunc-estimated CSS, then the MAG’s extraneous sequence content likely belongs to the same or a related taxon but still contains extra protein. The MAG may be a chimera of multiple highly related species or strains. Conversely, if a MAG is low in CheckM2-estimated contamination but high in gunc-estimated CSS, the extraneous sequence content may be non-redundant proteins from more distantly related taxa that nevertheless aligns to the reference genome (e.g. different genera). At least for this pilot dataset, there seems to be a stronger correlation between the CSS and the proportion unaligned bases than CheckM2-estimated contamination (Figure 6B, C), potentially indicating that there is more inter-species chimerism within a MAG. Both metrics only consider the over- or under-abundance of protein content. If there are contaminant non-protein-coding sequences, or sufficient synonymous mutation-based variation, that unalignable sequence material would not be considered by either metric. In contrast, strain heterogeneity-which measures the proportion of multiple copies of single-copy genes with high minimum amino acid similarity-naturally does not correllate with unaligned material, which by definition does not pass a sequence similarity threshold (Figure 6D).

**Fig. 6.**
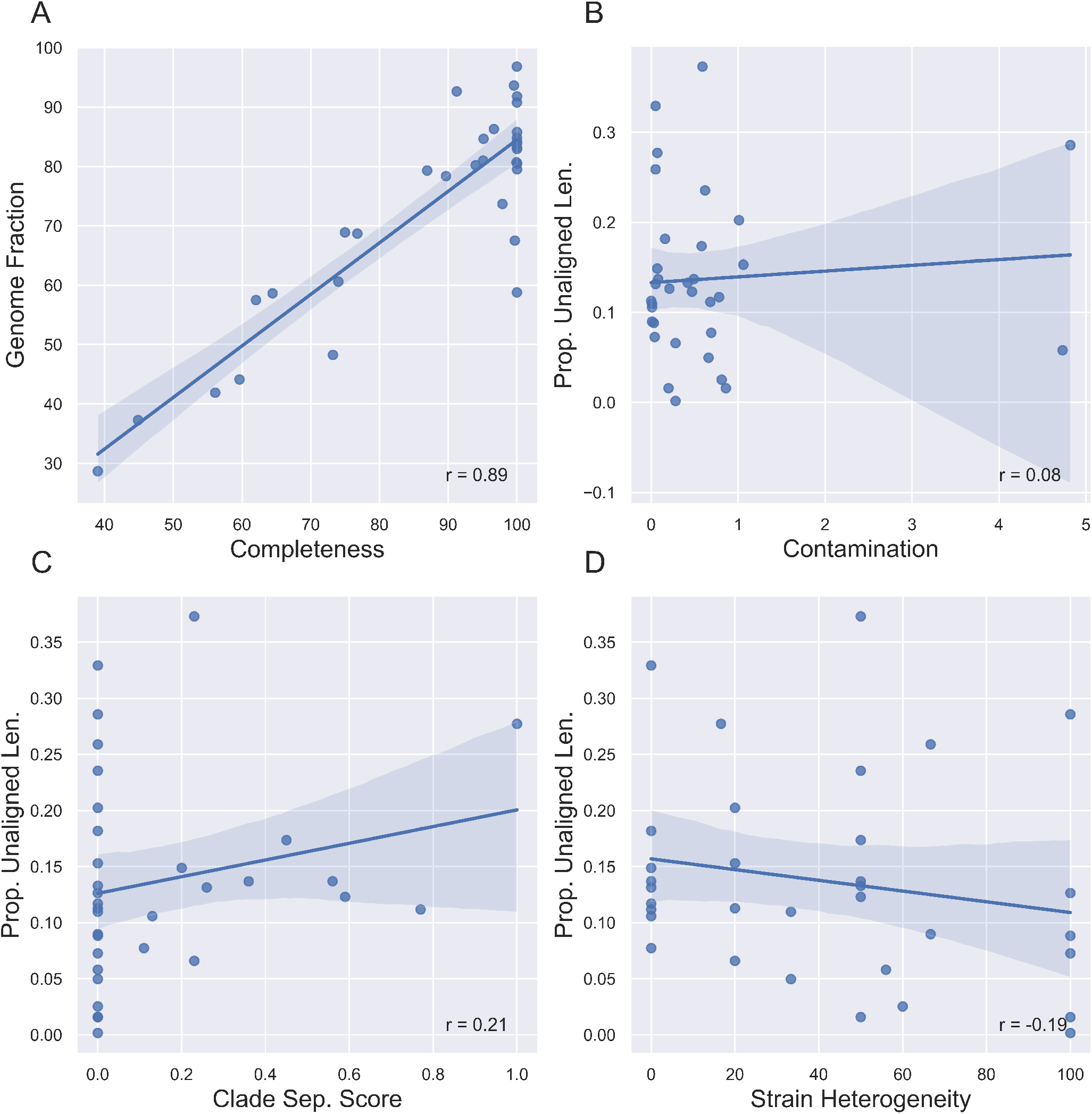
Reference-free metrics are sometimes correlated with reference-based metrics. (A) Completeness is highly correlated with QUAST-estimated genome fraction (the fraction of the reference genome that the MAG aligned to). (B, C) Contamination and clade separation score (CSS) do not correlate with the proportion of unaligned sequence material. (D) Strain heterogeneity is mildly inversely correlated with the proportion of unaligned material.

Consistently, a larger proportion of the MAG is alignable to the reference genome it is classified as, rather than the converse (Supp. Figure 2). Much of the reference genome material is missing in the MAG, which is corroborated by the median sizes of the MAGs-3 Mbp, which is small for a bacterial genome (Figure 4C, Figure 5J) [diCenzo and Finan, 2017]. Since the median average nucleotide identity between a MAG and the reference genome of its classified species (if available) was 97.34 (Figure 5G), the inferred MAGs are likely a specific substrain of the associated species. For example, the most abundant MAG in sample 4 was classified as *Cutibacterium acnes* and had an average coverage of 55.1X. 93.7% of the species reference genome was present in the MAG, and there were 573 positions with alternate alleles relative to the reference. Within the MAGs themselves, there was generally some degree of strain heterogeneity (Figure 5D).

The overall score is a summary statistic that takes into account gene content completeness, extra-lineage gene contamination, and overall sequence contiguity (2). While each MAG’s overall score is overall correlated with the MIMAG quality (Kruskal-Wallis H = 34.364, p = 3.45 *×* 10^−8^), it is surprisingly not correlated with whether or not the MAG is classified at the species level by GTDB-Tk (Kruskal-Wallis H = 0.0046, p = 0.946). The latter could be due to the fact that all MAGs were classified to the genus level and 75% of the MAGs were classifiable at the species level. Ergo, the overall score and MIMAG quality are high enough for both classified and unclassified that they are not likely to be distinguishing factors. Similarly, MIMAG quality is not correlated with species classification (*χ*-square = 3.822, p = 0.148). However, it is correlated with size (Kruskal-Wallis H = 7.291, p = 0.026) and even more strongly with N50 (Kruskal-Wallis H = 22.751, p = 1.147 *×* 10^−5^).

#### Comparing Minimum Contig Size Thresholds on MAG Quality

To determine if the minimum contig size threshold had appreciable impacts on MAG recovery and quality, we also binned Samples 3 and 7 (the smallest and largest short-read sequencing datasets, respectively) with a minimum size threshold of 1000 bp instead of 2500 bp. The results were highly dataset-dependent, and displayed no clear trends in terms of overall MAG quality (Supp. Figure 3 and Table 3). For maximally robust MAGs, it may be beneficial to bin assemblies with multiple minimum contig size thresholds, and then use an ensemble or dereplication tool like DAS Tool or dRep to find the most robust contig clusterings.

As expected, the sample 7 MAGs had lower CheckM2-estimated completeness and contamination when binned with a 2500 bp threshold than 1000 bp. Conversely, the sample 3 MAGs were higher in both quality metrics with a threshold of 2500 bp (Supp. Table 3). If the species-classified MAGs are considered alone, the average genome fraction (as measured by QUAST, represents the fraction of the associated reference genome found in the MAG) is higher with the 2500 bp cutoff, despite having the same average MAG sizes (Supp. Figure 3H, I). The MAGs assembled with the 1000 bp threshold also had proportionally more unaligned contigs and bases (Supp. Figure 3N, O).

In terms of recovered species, the same set of taxa were recovered in both 1000 and 2500 bp binning attempts of sample 3. The MAGs classified as *D. sp003523545* and *M. sp001878835* were larger with the 2500 bp threshold (Supp. Table 3).

However, the 1000 and 2500 bp attempts at MAG inference for sample 7 returned slightly different sets of taxa (Supp. Table 3). Specifically, the 2500 bp attempt at MAG inference did not recover *Chimaeribacter coloradensis* specifically, only a MAG that was classified as *Chimaeribacter* at the genus level (Supp. Table 3). The 1000 bp attempt at MAG inference recovered both the species *C. coloradensis* as well as a separate *Chimaeribacter*-only MAG, although the MAG was larger than the reference genome size (5.6 Mbp vs. 4.7 Mbp respectively), and the contamination and CSS were high (22.66, 0.52). The *Pseudomonas B*-classified MAG in the 2500 bp attempt was larger than the potential analog in the 1000 bp attempt, which was classified at a higher resolution as *P. B luteola*. However, the completeness of the 2500 bp attempt MAG was low, and contamination/clade separation score were high. As such, the MAG inferred with the 2500 bp size threshold is likely a chimera of several *Pseudomonas* species, as noted previously.

### Short-read Taxonomic Classification

The short-read taxonomic classification Jupyter notebook semi-automatically compares the 30 technical replicates (10 samples x 2 classifiers) using 2 alpha diversity metrics (taxonomic richness (i.e.: number of taxa and Shannon entropy) and 2 beta diversity metrics (Bray-Curtis dissimilarity and Jaccard distance) at different taxonomic ranks: phylum, class, order, family, genus, and species.

Despite discovering fewer unique taxa at every taxonomic rank overall, MetaPhlan4 almost always discovered more taxa than Kraken2-Bracken (Figure 7). When comparing shared taxa across all 20 technical replicates, both the Bray-Curtis and Jaccard distances between each replicate’s species profile indicates that there is almost total assortment of species by classifier (Figure 8 A,C). 137 unique species were discovered across both classifiers, w ith the vast majority (599/630, 95.1%) found by a single classifier. The vast majority-131/137 species (95.6%)-was discovered by only of the classifiers. Even for the 6/137 (4.4%) species discovered by both, the estimated relative abundance of the universally discovered species varied widely (Table 6). At the higher rank of phylum, there is less distinctive classifier-specific assortment, but the taxonomic profiles of the same sample generated by different classifiers tend not to cluster together (Figure 8B D). There are 3 phyla are present in a majority of classifier-sample replicates: Proteobacteria, Actinobacteria, and Firmicutes (Figure 9), which were previously noted as the most prevalent 3 phyla by the original MetaSUB publication [Danko et al., 2021]. The relative abundance accounted for by all of these phyla (per sample) are 0.941 (*±*0.081) ranging from [0.443, 1].

**Table 6.**
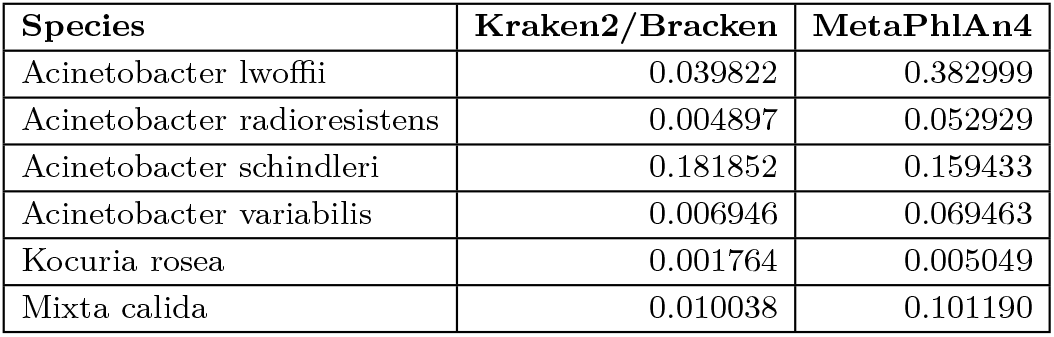
The relative abundance estimates for the 6 most high-abundance species discovered by Kraken2-Bracken and/or MetaPhlan4 can vary by a factor of 10.

**Fig. 7.**
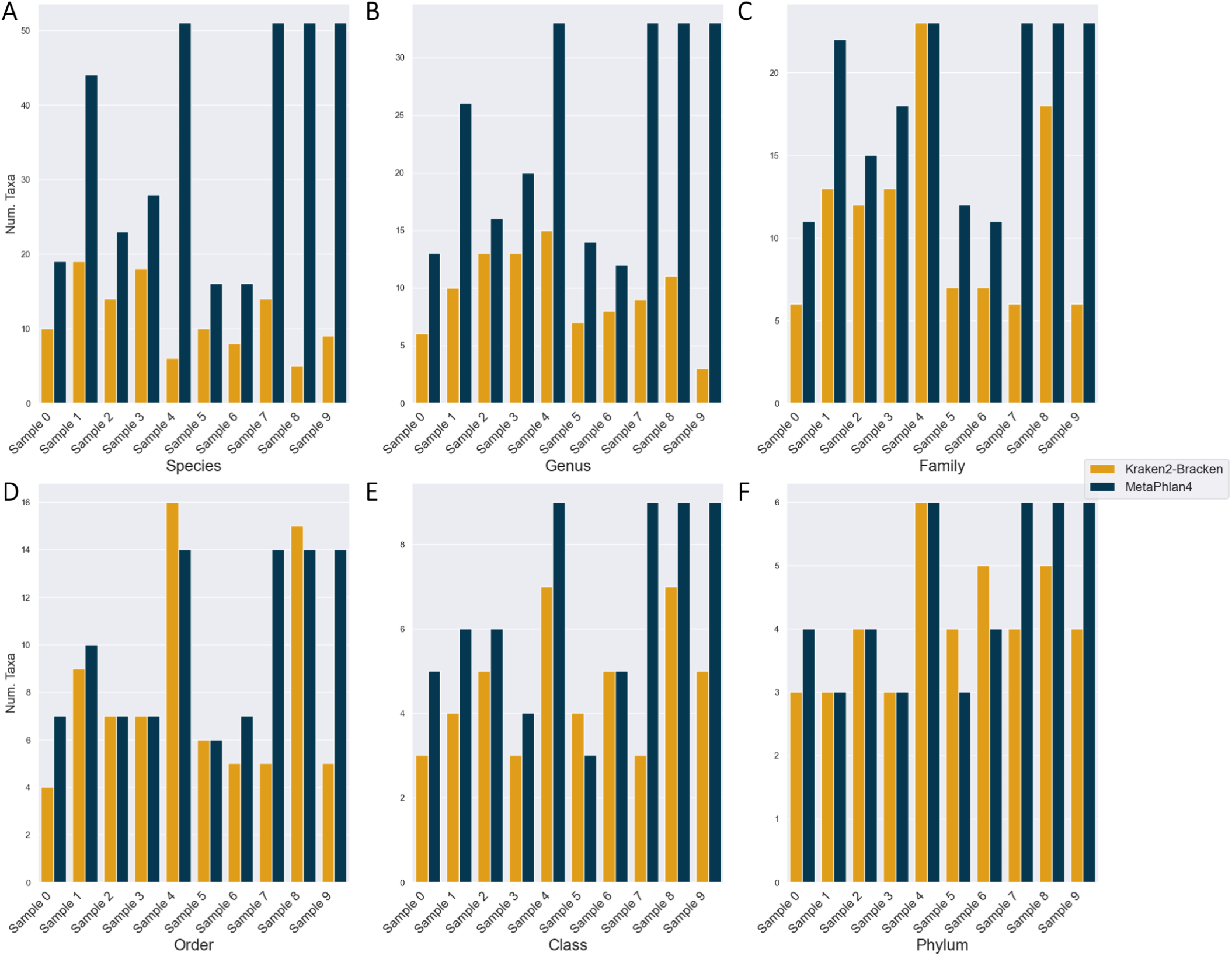
Each classifier detects different numbers of taxa across all ranks. (A) Species. (B) Genus. (C) Family. (D) Order. (E) Class. (F) Phylum.

**Fig. 8.**
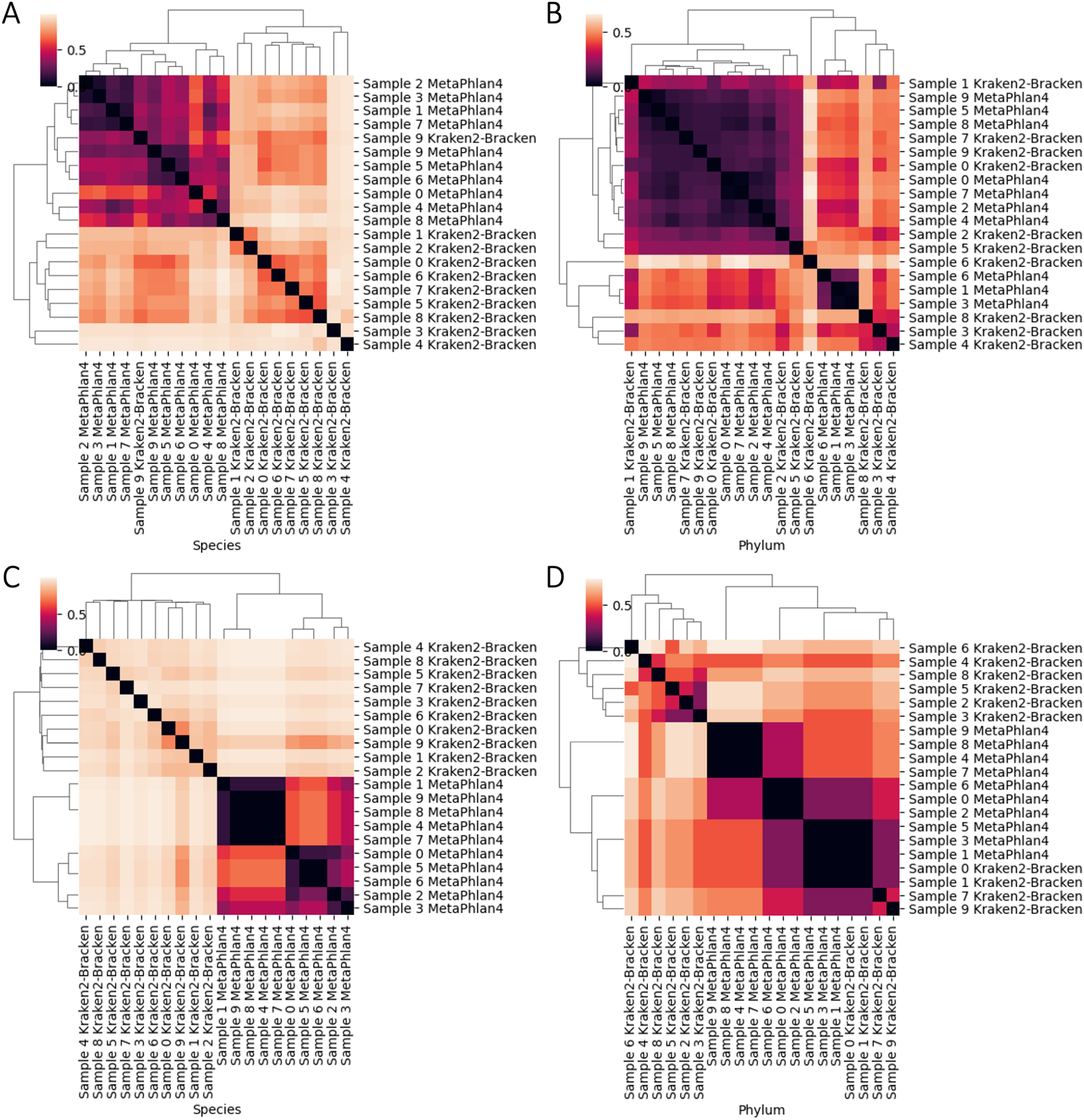
Taxonomic profiles are classifier-specific, not sample-specific, at the taxonomic rank of species but do not present any discernible patterns at the rank of phylum. Bray-Curtis and Jaccard distances between each replicate’s species (A, C) and phylum (C, D) profiles.

**Fig. 9.**
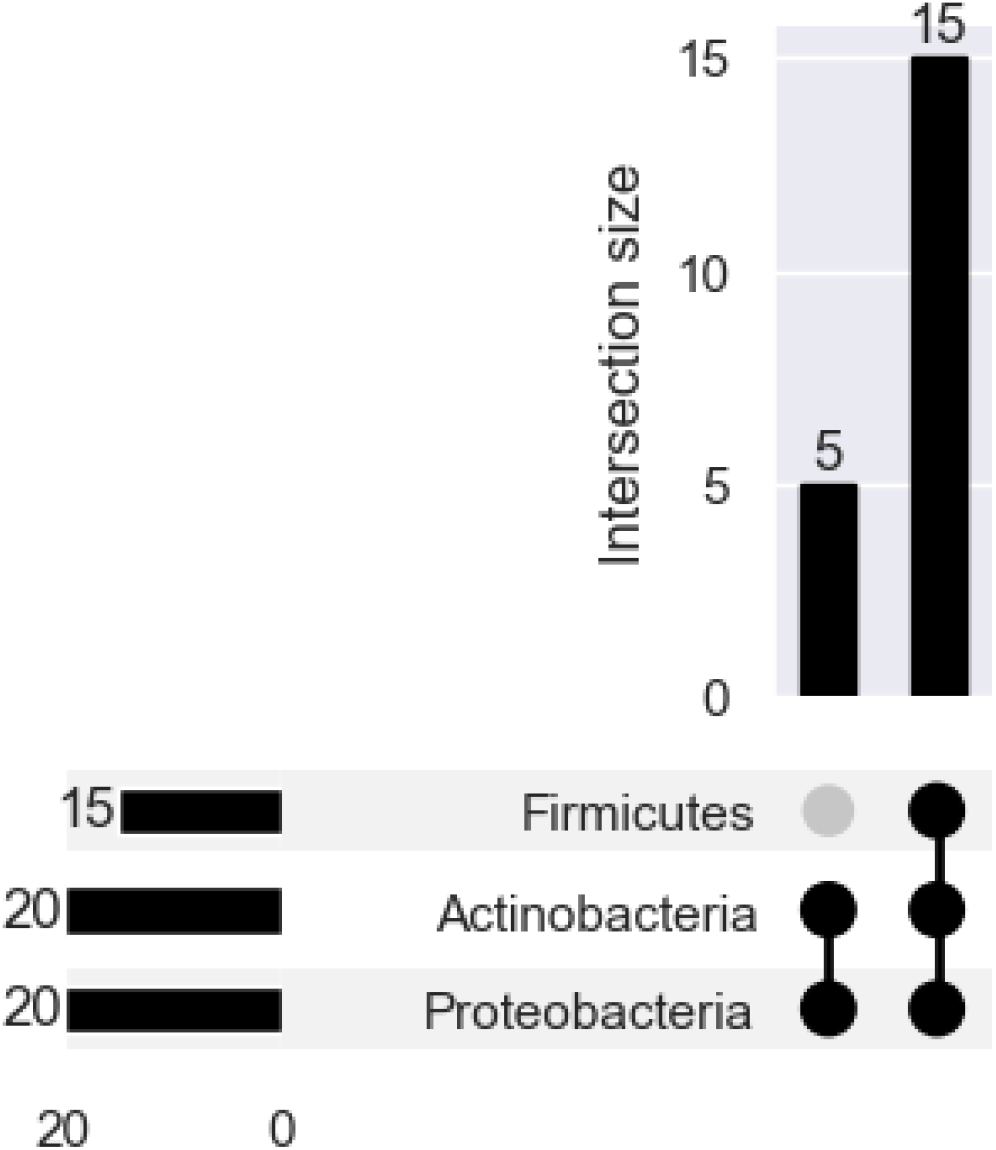
There are 3 phyla present in a majority or all of the 10 samples: Proteobacteria, Actinobacteria, Firmicutes, and Bacteroidota.

We conducted a small case study on the species discovered by the short-read classifiers in sample 7, and found that each classifier’s associated database contains some but not all of the species found in the others (Supplementary Materials). Unless custom databases are built from the same reference genome source, or multiple tools are used, it is difficult to compare classification strategies.

#### Comparing Short-Read Discovered and MAG Species

The number of unique species discovered across both short-read taxonomic classifiers far outnumbers the number of MAGs in each sample (Figure 7A vs. Table 5). If we consider a more stringent filter and only consider species that have been discovered in *≥* 10 classifier-sample replicates with a minimum relative abundance of at least 0.01, then the number of ‘verified’ s pecies i s much closer to the number of MAGs. However, the species discovered in by the short-read classifiers generally do not match the species classified by the M AGs (Table 7). By virtue of how ‘verified’ was defined, the 16 species are common to all 10 samples-6 having been discovered by both Kraken2-Bracken and MetaPhlan4, and 10 having been discovered by only Kraken2-Bracken in all 10 samples. When comparing highly prevalent short-read-discovered species to inferred MAGs, there is in fact more diversity in the number of MAG species discovered across all 10 samples (30) as well as more diversity between samples (Table 7). Of the species in Table 7, there are 9 ‘verified’ short-read and 17 M AG s pecies that d o n ot appear i n the list of 75 most abundant taxa (i.e.: the ‘core’ and part of the ‘sub-core’ global urban microbiome) in [Danko et al., 2021]. The most abundant bacterial species from the original study-*Cutibacterium acnes*-appears in the 3/10 samples (Table 7) [Danko et al., 2021].

**Table 7.**
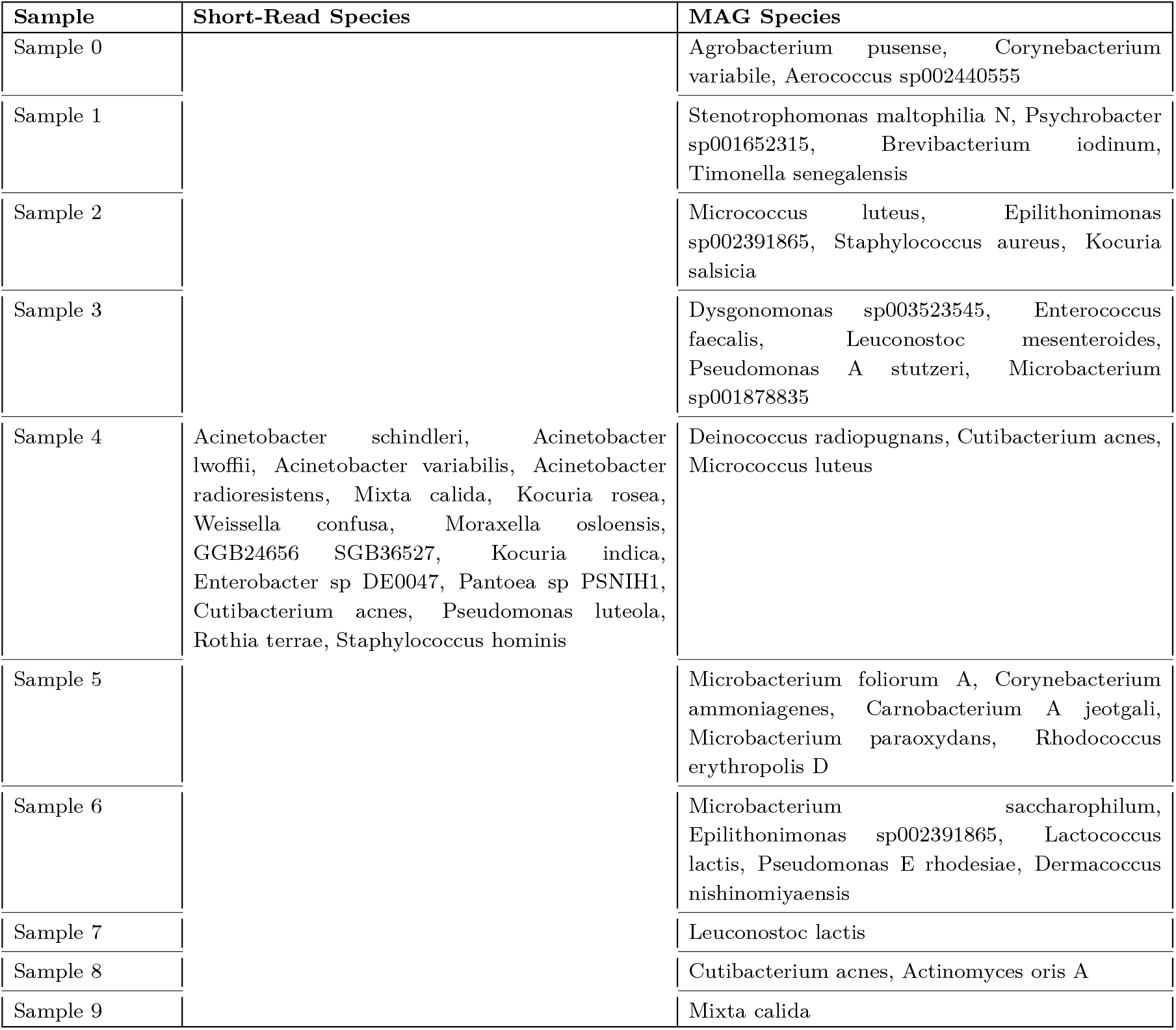
Most ‘verified’ (i.e.: discovered in at least 10 classifier-sample replicates, minimum relative abundance of 0.01) short-read species do not overlap with MAG species, and vice versa.

While there are likely more species per sample than binning algorithms are inferring, it is likely that the undiscovered species were not complete or stable enough for DAS Tool to confidently associate sets of contigs with each other.

### Virus and Phage Inference

A small fraction of the total number of contigs are flagged by the virus/phage-inference algorithms as potentially containing viral or phage sequence matter (Supp. Table 6). However, of those contigs, only around 10% of the flagged contigs are actually classifiable at the species i.e.: align to a virus or phage reference genome in RefSeq.

The largest number of virus/phage-flagged contigs was found in sample 7, which had the largest number of reads and assembly in terms of bases (Supp. Table 6). CheckV quality does not map well to RefSeq classifiability; only 17/26, 65.4% of CheckV-assessed complete viruses were species-classifiable. At best, the average CheckV-assessed completeness was on average 38.2% in sample 3, which contained 2044 algorithm flagged contigs (the second-most, next to sample 7) (Supp. Table 7).

There are 332 unique species-classifiable viruses, with the largest number in the sample with the largest number of reads, assembled bases, flagged contigs, and Blast-aligned contigs Of these, 259/332 (78.0%) are unique to a single sample. The same virus or phage species was typically discovered multiple times in the same sample (Table 8). Taking the low completeness per contig into account, each contig could either be fragments of the same isolate, or there could be viral/phage strain diversity in the sample despite the low (‘verified’) bacterial diversity. 11/332 (3.3%) of species were found in 4+ samples (Figure 10). Of these, 10/11 (90.9%) aligned to bacteriophage reference genomes.

**Table 8.** Each virus or phage species discovered in a sample is represented by on average *>* 1 contig in that sample.

| Sample | Num. Viruses/Phages | Avg. Num. Contigs |
| --- | --- | --- |
| Sample 0 | 15 | 1.40 |
| Sample 1 | 45 | 1.71 |
| Sample 2 | 60 | 1.82 |
| Sample 3 | 61 | 3.64 |
| Sample 4 | 36 | 1.22 |
| Sample 5 | 16 | 1.63 |
| Sample 6 | 37 | 2.95 |
| Sample 7 | 130 | 3.32 |
| Sample 8 | 8 | 1.38 |
| Sample 9 | 37 | 2.19 |

**Fig. 10.**
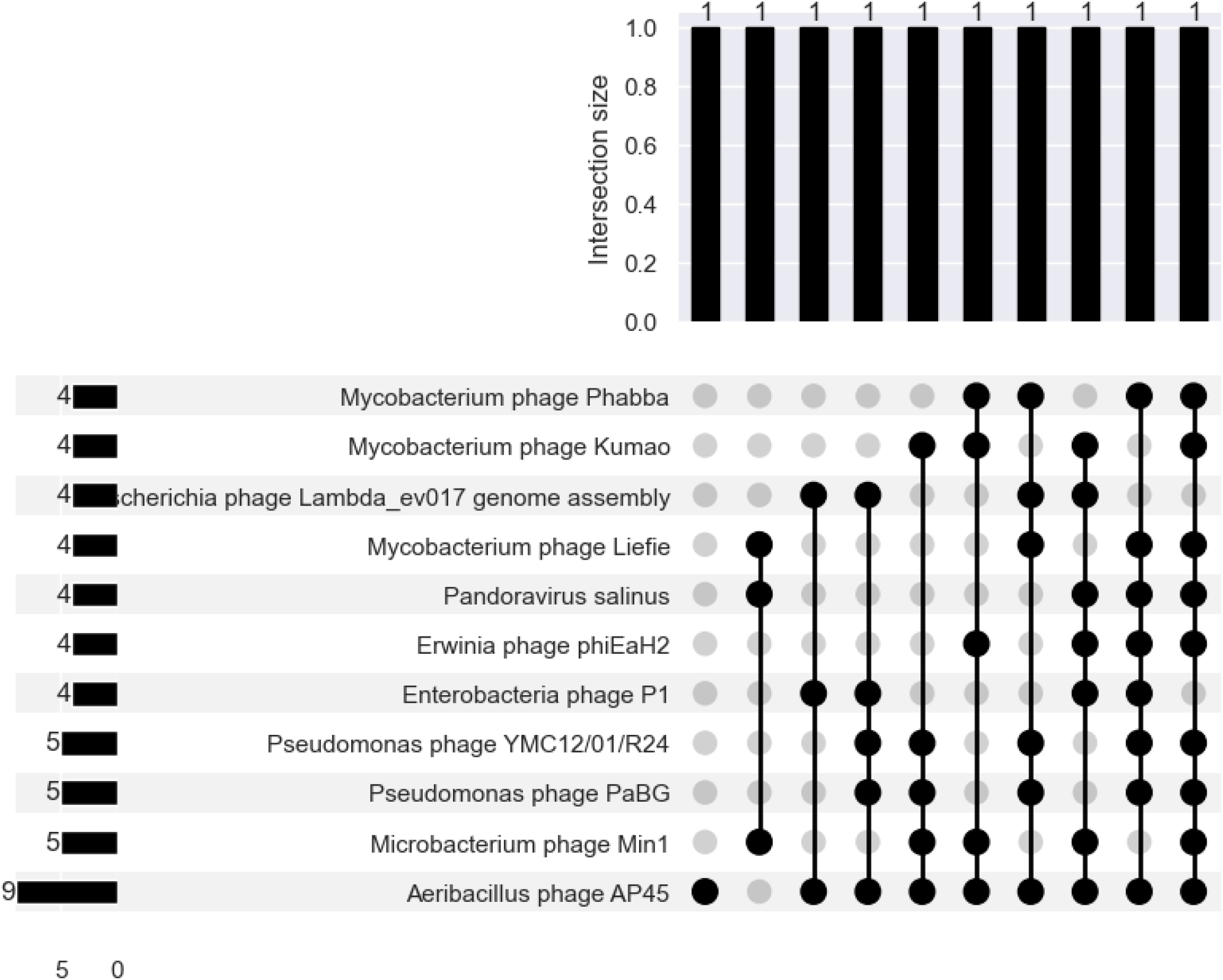
There is more overlap between the virus and/or phage profiles of different samples than there are bacteria.

#### Comparing Virus-Phage and MAG Classifications

To maintain consistency from the short-read taxonomy section, we selected sample 4 for a case study. When we cross-referenced algorithm-flagged contigs with contig assignments to MAGs, we found that 20/44 (45.5%) candidate virus/phage contigs are found in MAGs. Only 1/44 (2.3%) contigs is labeled as a provirus by CheckV-Bixzunavirus I3. The contig is associated with a MAG, but MAG was classified as *Micrococcus luteus* and is not of the genus Mycobacterium, which is the phages’ known host [Reddy and Gopinathan, 1987].

As with the above example, most of the phage-MAG associations are not within the narrow host range of phages. Most phages can only infect several strains of bacteria from the same species, though there are documented cases of phages infecting bacteria from different genera [Ross et al., 2016]. Of the non-phage contigs (8/44, 18.2%), none are associated with MAGs except 3/4 Pandoraviruses, which are oddly associated with Micrococcus luteus.

### Gene Cataloguing

There are 407, 098 individual gene annotations found across all contigs across all samples with an average of 3.974 *±*28.98 (range: [1, 3642]). The median number of genes annotated per contig is 1, which agrees with the shortness of most contigs (per-sample averages are 262 - 930 bp). The vast majority of contigs are extremely short->99.5% are <2500 bp.

The vast majority of genes cannot be assigned to a cluster of orthologous genes (COG), which are families of orthologous protein-coding genes (Supp. Table 8). Generally, the larger the dataset and/or assembly, the more COG representatives are found in the assembly. Samples 4, 7, and 8 do not have gene annotations because the associated Bakta jobs did not complete after a week of running the bakta command, likely because the assemblies were too large to completely annotate. Most genes, if they are assigned to a COG, belong to the metabolism category (Figure 11A). However, the COG with the largest number of genes is that of translation and ribosomal structure (Figure 11C). There were very few genes annotated in energy production and conversion, extracellular structures, and cytoskeleton COGs (Figure 11). It is unknown whether urban microbiome assemblies simply contain very few genes of those functions, or are they difficult to assemble or annotate (e.g. not conserved enough or difficult to find via HMM). There were not many cell motility-annotated genes either, which is interesting given one would expect to find many free-living species in the urban milieu.

**Fig. 11.**
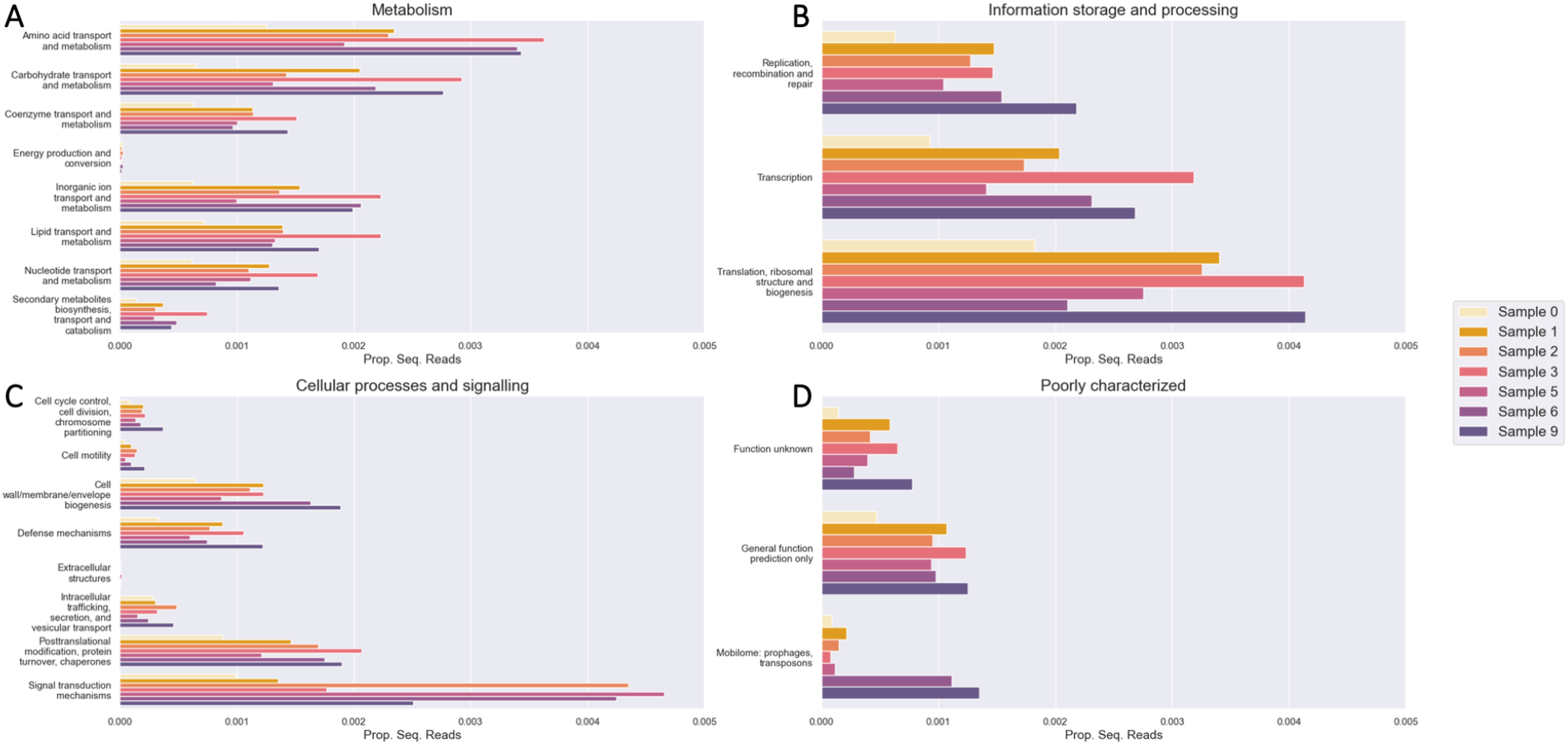
(A) Metabolism, (B) information and storage, (C) cellular process and signaling, and (D) poorly characterized clusters of orthologous genes (COGs) annotated in each sample. The x-axis refers to the proportion of sequencing reads aligned to genes from a COG.

### Computational Resources Used

If executed as an end-to-end workflow, all modules in our pipeline can be used to efficiently process sequencing data on a high-performance computing (HPC) cluster. The workflow was designed to optimize resource utilization and minimize processing time (Table 9). Our configuration allowed us to process large-scale metagenomic datasets efficiently, enabling comprehensive analysis of microbial communities across multiple samples. The modular nature of our workflow allows for scalability and adaptability to different computational environments, from local workstations to cloud-based infrastructures.

**Table 9.**
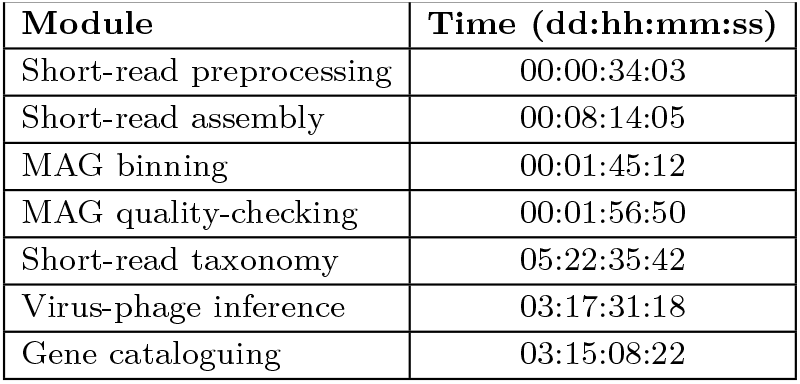
Time required to process all 10 samples with 64 CPUs available to the Snakemake command.

| Module | Time (dd:hh:mm:ss) |
| --- | --- |
| Short-read preprocessing | 00:00:34:03 |
| Short-read assembly | 00:08:14:05 |
| MAG binning | 00:01:45:12 |
| MAG quality-checking | 00:01:56:50 |
| Short-read taxonomy | 05:22:35:42 |
| Virus-phage inference | 03:17:31:18 |
| Gene cataloguing | 03:15:08:22 |

## Discussion

In this paper, we showcase CAMP, a fully modular system comprised of core metagenomic analysis tools, and demonstrated its utility in current and future metagenomic research through a pilot-size case study with real metagenomics data. CAMP is by no means “yet another metagenomic pipeline”, but a future-facing infrastructure built with the hope of laying the foundation for future metagenomic research. It is thoughtfully designed with a comprehensive set of tools covering almost all aspects of core metagenomics analyses, from preprocessing to functional inference, with the support for both short and long reads. To the best of our knowledge, CAMP is the most modular and functionally comprehensive metagenomic analysis system available to date, allowing researchers the maximum flexibility to design customized workflows catering to their specific research needs. In addition to user experience, CAMP has been designed with developer accessibility in mind, with a standard module structure setup for easy development of new modules using the templates provided.

While other pipelines may be sufficient for small-scale projects, CAMP is irreplaceable in many potential large-scale exploratory future projects. For example, the modular sandbox design makes CAMP an ideal candidate for module-wise tool benchmarking. Considering the amount of metagenomic analysis tools available and the variability of results produced by different tools, the amount of literature on metagenomic tool benchmarking is surprisingly low. This can be attributed to a combination of factors such as the lack of gold-standard datasets and the computational load. Despite being a current challenge, tool benchmarking is indispensable and unavoidable in the future as better quality datasets continue to emerge as a result of advancement in biotechnology and our understanding. With its modularity and HPC-compatibility, CAMP is prepared to address such needs whenever they arise in the future. Its modular design with standardized input and output formats allow convenient distribution of workloads and reproducibility of results in large-scale collaborative efforts. In addition to the modules mentioned in this paper, there are many more CAMP modules currently under active development, including long-read taxonomic profiling and decontamination. As our next milestone, we will further expand the utilities of CAMP to include multi-sample co-assembly and co-binning, variant and strain analysis, and metatranscriptomics. To further enhance reproducibility and interoperability for users with root access, we also plan to offer a containerized implementation of CAMP using Singularity.

We note that there is not a single tool or toolkit that is sufficient to meet every single need. With greater flexibility of CAMP comes greater responsibility on the side of the researchers. With less hard-coded defaults in parameters, CAMP places more weight on the expertise and discretion of the researchers in experiment design and parameter selection. The soft pauses and semi-guided visualizations at the end of each module also provide users the opportunity to spot any potential inappropriate protocols and make modifications promptly.

We envision CAMP not to be another short-lived pipeline, but a sustainable ecosystem of high-quality toolkit for all core aspects of metagenomic analysis that will be consistently expanded, maintained, and fortified to lay a solid foundation for future metagenomic research.

## Author Contributions Statement

L.M. and B.T. jointly conceived of the CAMP modular analysis concept, with design contributions from D.D. L.M., B.T., I.H., and C.M. jointly managed project development. L.M. developed the modular template and short-read assembly module, with contributions from B.T, C.R., and C.F. L.M., B.T., and C.F. developed the short-read quality-control module with contributions from C.R. L.M. developed the MAG binning and quality-control modules. B.T., L.M., and C.F. developed the short-read taxonomy module with contributions from S.Z. and J.H. B.T. and L.M. developed the gene cataloguing module. R.B.T. developed the viral inference module with contributions from J.S.A. and L.M. M.T. and B. Turhan developed the Nanopore long-read quality control module with contributions from L.M. and D.M. C.R. developed the decontamination module with contributions from M.K., A.N., Z.I., S.K., and N.N. D.H., V.K., A.F., and A.K. developed the mOTUs module. W.W. implemented parts of all modules. L.M. did base-case testing for all of the modules. A.H.S., J.C.S., and A.G.L. contributed to the debugging process. L.M. wrote the manuscript with suggestions from all of the other authors. W.W., A.G.L. and E.E. contributed to the editing process. B.T. created the ReadTheDocs.

## Acknowledgments

We thank Haruo Suzuki and Yinka Osuolale for their copy-editing suggestions. We gratefully acknowledge Poland’s high-performance Infrastructure PLGrid (HPC Centers: ACK Cyfronet AGH, PCSS, CI TASK, WCSS) for providing computing facilities.

## Data Availability Statement

The raw sequencing sequencing data can be found under the SRA accession ID PRJNA732392, as well as the GeoSeeq repository.

## Box 1: Standardized module architecture for system-wide navigability and extendability

`workflow/`: Contains all of the module’s functional code and scripts from external tools that are under an open-source license.

`module_slug.py`: The Click-based command-line interface that wraps a Snakemake-based workflow.

`Snakefile`: Workflow that itself wraps Snakemake rules, which call external algorithms or internal functions (utils.py) to process data.

`utils.py`: Internal functions such as dataset ingestion and working directory setup.

`ext/`: External scripts and smaller data files required by algorithms used in the workflow.

`configs/`: Contains all settings files.

`parameter.yaml and resource.yaml`: Instead of relying on Snakemake’s dictionary-based system of specifying parameters and resources on the command line,algorithm parameters and computational resource allocations have been unified into single YAML files. Users can: i) use the default YAMLs provided, ii) modify the values in the YAMLs according to their dataset properties, or iii) make multiple copies of the YAMLs to analyze datasets of different sizes and biological origins.

`samples.csv`: A standardized input file where each row corresponds to a sample. The first column is always the sample’s name, and every column thereafter contains paths to that sample’s data.

`conda/`: Conda environment recipes. The main environment is always called `module_slug.yaml` and should be set up immediately after cloning the module to a local directory. All other Conda environments will be set up the first time the workflow is run into a directory called `conda_envs/` in the module’s top-level directory.

`sbatch/`: A Snakemake profile that enables the module’s rules to be run using the Slurm job scheduler.

`test_data/`: Test data to ensure proper module setup, and an example work directory to demonstrate its structure, and input, intermediate, and output file formats.

`dataviz.ipynb`: Guided and semi-automated data visualization of the module’s final output using Jupyter notebook. Users can use the built-in graphing functions as is or customize as needed.

## References

Alexandre Almeida, Alex L. Mitchell, Miguel Boland, Samuel C. Forster, Gregory B. Gloor, Aleksandra Tarkowska, Trevor D. Lawley, and Robert D. Finn. A new genomic blueprint of the human gut microbiota. Nature, 568(7753): 499–504, 2019. doi: 10.1038/s41586-019-0965-1.

Johannes Alneberg, Brynjar Smári Bjarnason, Ino De Bruijn, Melanie Schirmer, Joshua Quick, Umer Z. Ijaz, Leo Lahti, Nicholas J. Loman, Anders F. Andersson, and Christopher Quince. Binning metagenomic contigs by coverage and composition. Nature Methods, 11(11):1144–1146, oct 2014. ISSN 15487105. doi: 10.1038/nmeth.3103. URL https://github.com/.

S. Andrews. A quality control tool for high throughput sequence data, 2010. URL http://www.bioinformatics.babraham.ac.uk/projects/fastqc/.

Dmitry Antipov, Anton Korobeynikov, Jeffrey S. McLean, and Pavel A. Pevzner. HybridSPAdes: An algorithm for hybrid assembly of short and long reads. Bioinformatics, 32(7):1009–1015, 2016. ISSN 14602059. doi: 10.1093/bioinformatics/btv688.

Dmitry Antipov, Mikhail Raiko, Alla Lapidus, and Pavel A. Pevzner. Metaviral SPAdes: Assembly of viruses from metagenomic data. Bioinformatics, 36(14): 4126–4129, 2020. ISSN 14602059. doi: 10.1093/bioinformatics/btaa490.

Mohammad Bahram, Tarquin Netherway, Clémence Frioux, Pamela Ferretti, Luis Pedro Coelho, Stefan Geisen, Peer Bork, and Falk Hildebrand. Metagenomic assessment of the global diversity and distribution of bacteria and fungi. Environmental Microbiology, 23(1): 316–326, 2021.

Aitor Blanco-Miguez, Francesco Beghini, Fabio Cumbo, Lauren J. McIver, Kelsey N. Thompson, Moreno Zolfo, Paolo Manghi, Leonard Dubois, Kun D. Huang, Andrew Maltez Thomas, and et al. Extending and improving metagenomic taxonomic profiling with uncharacterized species with metaphlan 4. bioarXiv, 2022. doi: 10.1101/2022.08.22.504593.

George Bouras, Louise M. Judd, Robert A. Edwards, Sarah Vreugde, Timothy P. Stinear, and Ryan R. Wick. How low can you go? Short-read polishing of Oxford Nanopore bacterial genome assemblies. Microbial Genomics, 10(6): 1–9, 2024. ISSN 20575858. doi: 10.1099/mgen.0.001254.

Robert M Bowers, Nikos C Kyrpides, Ramunas Stepanauskas, Miranda Harmon-Smith, Devin Doud, T B Reddy, Frederik Schulz, Jessica Jarett, Adam R Rivers, Emiley A Eloe-Fadrosh, and et al. Minimum information about a single amplified genome (misag) and a metagenome-assembled genome (mimag) of bacteria and archaea. Nature Biotechnology, 35(8): 725–731, 2017. doi: 10.1038/nbt.3893.

Ilana L. Brito, Thomas Gurry, Shijie Zhao, Katherine Huang, Sarah K. Young, Terrence P. Shea, Waisea Naisilisili, Aaron P. Jenkins, Stacy D. Jupiter, Dirk Gevers, and et al. Transmission of human-associated microbiota along family and social networks. Nature Microbiology, 2019. doi: 10.1101/540252.

Benjamin Buchfink, Chao Xie, and Daniel H. Huson. Fast and sensitive protein alignment using DIAMOND. Nature Methods, 12(1): 59–60, 2014. ISSN 15487105. doi: 10.1038/nmeth.3176.

Brian Bushnell. BBMap: a fast, accurate, splice-aware aligner. Lawrence Berkeley National Laboratory, pages LBNL Report : LBNL–7065E, 2014. URL https://escholarship.org/uc/item/1h3515gn.

Antonio Pedro Camargo, Simon Roux, Frederik Schulz, Michal Babinski, Yan Xu, Bin Hu, Patrick S.G. Chain, Stephen Nayfach, and Nikos C. Kyrpides. Identification of mobile genetic elements with geNomad. Nature Biotechnology, 42(8): 1303–1312, 2024. ISSN 15461696. doi: 10.1038/s41587-023-01953-y. URL http://dx.doi.org/10.1038/s41587-023-01953-y.

Diego D Cambuy, Felipe H Coutinho, and Bas E Dutilh. Contig annotation tool cat robustly classifies assembled metagenomic contigs and long sequences. BioRxiv, page 072868, 2016.

Pierre-Alain Chaumeil, Aaron J Mussig, Philip Hugenholtz, and Donovan H Parks. Gtdb-tk: A toolkit to classify genomes with the genome taxonomy database. Bioinformatics, 2019. doi: 10.1093/bioinformatics/btz848.

Lin-Xing Chen, Karthik Anantharaman, Alon Shaiber, A. Murat Eren, and Jillian F. Banfield. Accurate and complete genomes from metagenomes. Genome Research, 2019. doi: 10.1101/808410.

Shifu Chen, Yanqing Zhou, Yaru Chen, and Jia Gu. Fastp: An ultra-fast all-in-one FASTQ preprocessor. Bioinformatics, 34(17): i884–i890, 2018. ISSN 14602059. doi: 10.1093/bioinformatics/bty560.

Alex Chklovski, Donovan H. Parks, Ben J. Woodcroft, and Gene W. Tyson. CheckM2: a rapid, scalable and accurate tool for assessing microbial genome quality using machine learning. Nature Methods, 20(8): 1203–1212, 2023. ISSN 15487105. doi: 10.1038/s41592-023-01940-w.

conda contributors. conda: A system-level, binary package and environment manager running on all major operating systems and platforms. URL https://github.com/conda/conda.

Cookiecutter. Cookiecutter/cookiecutter: A cross-platform command-line utility that creates projects from cookiecutters (project templates), 2022. URL https://github.com/cookiecutter/cookiecutter.

Wouter De Coster and Rosa Rademakers. Sequence analysis NanoPack2 : population-scale evaluation of long-read sequencing data. Bioinformatics, 39(May):0–2, 2023.

David Danko, Daniela Bezdan, Evan E. Afshin, Sofia Ahsanuddin, Chandrima Bhattacharya, Daniel J. Butler, Kern Rei Chng, Daisy Donnellan, Jochen Hecht, Katelyn Jackson, and et al. A global metagenomic map of urban microbiomes and antimicrobial resistance. Cell, 184(13), 2021. doi: 10.1016/j.cell.2021.05.002.

Nicole M. Davis, DIana M. Proctor, Susan P. Holmes, David A. Relman, and Benjamin J. Callahan. Simple statistical identification and removal of contaminant sequences in marker-gene and metagenomics data. Microbiome, 6(1): 1–14, 2018. ISSN 20492618. doi: 10.1186/s40168-018-0605-2.

Paolo Di Tommaso, Maria Chatzou, Evan W Floden, Pablo Prieto Barja, Emilio Palumbo, and Cedric Notredame. Nextflow enables reproducible computational workflows. Nature Biotechnology, 35 (4):316–319, 2017. doi: 10.1038/nbt.3820.

New Text Compilation4George C. diCenzo and Turlough M. Finan. The divided bacterial genome: Structure, function, and evolution. Microbiology and Molecular Biology Reviews : MMBR, 81(3): e00019–17, August 2017. ISSN 1092-2172. doi: 10.1128/MMBR.00019-17. URL https://www.ncbi.nlm.nih.gov/pmc/articles/PMC5584315/.

Marine Djaffardjy, George Marchment, Clémence Sebe, Raphaël Blanchet, Khalid Belhajjame, Alban Gaignard, Frédéric Lemoine, and Sarah Cohen-Boulakia. Developing and reusing bioinformatics data analysis pipelines using scientific workflow systems. Computational and Structural Biotechnology Journal, 21: 2075–2085, 2023. ISSN 20010370. doi: 10.1016/j.csbj.2023.03.003. URL https://linkinghub.elsevier.com/retrieve/pii/S2001037023001010.

Philip Ewels, Måns Magnusson, Sverker Lundin, and Max Käller. Multiqc: Summarize analysis results for multiple tools and samples in a single report. Bioinformatics, 32(19): 3047–3048, 2016. doi: 10.1093/bioinformatics/btw354.

Philip A. Ewels, Alexander Peltzer, Sven Fillinger, Harshil Patel, Johannes Alneberg, Andreas Wilm, Maxime Ulysse Garcia, Paolo Di Tommaso, and Sven Nahnsen. The nf-core framework for community-curated bioinformatics pipelines. Nature Biotechnology, 38(3):276–278, March 2020. ISSN 1087-0156, 1546-1696. doi: 10.1038/s41587-020-0439-x. URL https://www.nature.com/articles/s41587-020-0439-x.

GabeAl. Gabeal/utree: K-mer searching with trees, 2022. URL https://github.com/GabeAl/UTree.

Dirk Gevers, Rob Knight, Joseph F Petrosino, Katherine Huang, Amy L McGuire, Bruce W Birren, Karen E Nelson, Owen White, Barbara A Methé, and Curtis Huttenhower. The human microbiome project: a community resource for the healthy human microbiome. 2012.

Jiarong Guo, Ben Bolduc, Ahmed A. Zayed, Arvind Varsani, Guillermo Dominguez-Huerta, Tom O. Delmont, Akbar Adjie Pratama, M. Consuelo Gazitúa, Dean Vik, Matthew B. Sullivan, and Simon Roux. VirSorter2: a multi-classifier, expert-guided approach to detect diverse DNA and RNA viruses. Microbiome, 9 (1):1–13, 2021. ISSN 20492618. doi: 10.1186/s40168-020-00990-y.

Tatiana A. Gurbich, Alexandre Almeida, Martin Beracochea, Tony Burdett, Josephine Burgin, Guy Cochrane, Shriya Raj, Lorna Richardson, Alexander B. Rogers, Ekaterina Sakharova, Gustavo A. Salazar, and Robert D. Finn. Mgnify genomes: A resource for biome-specific microbial genome catalogues. Journal of Molecular Biology, 435(14):168016, July 2023. ISSN 00222836. doi: 10.1016/j.jmb.2023.168016. URL https://linkinghub.elsevier.com/retrieve/pii/S0022283623000724.

Alexey Gurevich, Vladislav Saveliev, Nikolay Vyahhi, and Glenn Tesler. QUAST: Quality assessment tool for genome assemblies. Bioinformatics, 29(8): 1072–1075, 2013. ISSN 13674803. doi: 10.1093/bioinformatics/btt086.

Kasper D. Hansen, Steven E. Brenner, and Sandrine Dudoit. Biases in illumina transcriptome sequencing caused by random hexamer priming. Nucleic Acids Research, 38(12):e131–e131, July 2010. ISSN 1362-4962, 0305-1048. doi: 10.1093/nar/gkq224. URL https://academic.oup.com/nar/article-lookup/doi/10.1093/nar/gkq224.

Maria Hauser, Martin Steinegger, and Johannes Söding. MMseqs software suite for fast and deep clustering and searching of large protein sequence sets. Bioinformatics, 32(9): 1323–1330, 2016. ISSN 14602059. doi: 10.1093/bioinformatics/btw006.

Tom CJ Hill, Kerry A Walsh, James A Harris, and Bruce F Moffett. Using ecological diversity measures with bacterial communities. FEMS microbiology ecology, 43(1): 1–11, 2003.

Steven Hofmeyr, Rob Egan, Evangelos Georganas, Alex C. Copeland, Robert Riley, Alicia Clum, Emiley Eloe-Fadrosh, Simon Roux, Eugene Goltsman, Aydin Buluç, and et al. Terabase-scale metagenome coassembly with metahipmer. Scientific Reports, 10(1), 2020. doi: 10.1038/s41598-020-67416-5.

Laura A Hug, Brett J Baker, Karthik Anantharaman, Christopher T Brown, Alexander J Probst, Cindy J Castelle, Cristina N Butterfield, Alex W Hernsdorf, Yuki Amano, Kotaro Ise, et al. A new view of the tree of life. Nature microbiology, 1(5): 1–6, 2016.

Curtis Huttenhower, Dirk Gevers, Rob Knight, Sahar Abubucker, Jonathan H. Badger, Asif T. Chinwalla, Heather H. Creasy, Ashlee M. Earl, Michael G. Fitzgerald, Robert S. Fulton, Michelle G. Giglio, Kymberlie Hallsworth-Pepin, Elizabeth A. Lobos, Ramana Madupu, Vincent Magrini, John C. Martin, Makedonka Mitreva, Donna M. Muzny, Erica J. Sodergren, James Versalovic, Aye M. Wollam, Kim C. Worley, Jennifer R. Wortman, Sarah K. Young, Qiandong Zeng, Kjersti M. Aagaard, Olukemi O. Abolude, Emma Allen-Vercoe, Eric J. Alm, Lucia Alvarado, Gary L. Andersen, Scott Anderson, Elizabeth Appelbaum, Harindra M. Arachchi, Gary Armitage, Cesar A. Arze, Tulin Ayvaz, Carl C. Baker, Lisa Begg, Tsegahiwot Belachew, Veena Bhonagiri, Monika Bihan, Martin J. Blaser, Toby Bloom, Vivien Bonazzi, J. Paul Brooks, Gregory A. Buck, Christian J. Buhay, Dana A. Busam, Joseph L. Campbell, Shane R. Canon, Brandi L. Cantarel, Patrick S.G. Chain, I. Min A. Chen, Lei Chen, Shaila Chhibba, Ken Chu, Dawn M. Ciulla, Jose C. Clemente, Sandra W. Clifton, Sean Conlan, Jonathan Crabtree, Mary A. Cutting, Noam J. Davidovics, Catherine C. Davis, Todd Z. Desantis, Carolyn Deal, Kimberley D. Delehaunty, Floyd E. Dewhirst, Elena Deych, Yan Ding, David J. Dooling, Shannon P. Dugan, Wm Michael Dunne, A. Scott Durkin, Robert C. Edgar, Rachel L. Erlich, Candace N. Farmer, Ruth M. Farrell, Karoline Faust, Michael Feldgarden, Victor M. Felix, Sheila Fisher, Anthony A. Fodor, Larry J. Forney, Leslie Foster, Valentina Di Francesco, Jonathan Friedman, Dennis C. Friedrich, Catrina C. Fronick, Lucinda L. Fulton, Hongyu Gao, Nathalia Garcia, Georgia Giannoukos, Christina Giblin, Maria Y. Giovanni, Jonathan M. Goldberg, Johannes Goll, Antonio Gonzalez, Allison Griggs, Sharvari Gujja, Susan Kinder Haake, Brian J. Haas, Holli A. Hamilton, Emily L. Harris, Theresa A. Hepburn, Brandi Herter, Diane E. Hoffmann, Michael E. Holder, Clinton Howarth, Katherine H. Huang, Susan M. Huse, Jacques Izard, Janet K. Jansson, Huaiyang Jiang, Catherine Jordan, Vandita Joshi, James A. Katancik, Wendy A. Keitel, Scott T. Kelley, Cristyn Kells, Nicholas B. King, Dan Knights, Heidi H. Kong, Omry Koren, Sergey Koren, Karthik C. Kota, Christie L. Kovar, Nikos C. Kyrpides, Patricio S. La Rosa, Sandra L. Lee, Katherine P. Lemon, Niall Lennon, Cecil M. Lewis, Lora Lewis, Ruth E. Ley, Kelvin Li, Konstantinos Liolios, Bo Liu, Yue Liu, Chien Chi Lo, Catherine A. Lozupone, R. Dwayne Lunsford, Tessa Madden, Anup A. Mahurkar, Peter J. Mannon, Elaine R. Mardis, Victor M. Markowitz, Konstantinos Mavromatis, Jamison M. McCorrison, Daniel McDonald, Jean McEwen, Amy L. McGuire, Pamela McInnes, Teena Mehta, Kathie A. Mihindukulasuriya, Jason R. Miller, Patrick J. Minx, Irene Newsham, Chad Nusbaum, Michelle Oglaughlin, Joshua Orvis, Ioanna Pagani, Krishna Palaniappan, Shital M. Patel, Matthew Pearson, Jane Peterson, Mircea Podar, Craig Pohl, Katherine S. Pollard, Mihai Pop, Margaret E. Priest, Lita M. Proctor, Xiang Qin, Jeroen Raes, Jacques Ravel, Jeffrey G. Reid, Mina Rho, Rosamond Rhodes, Kevin P. Riehle, Maria C. Rivera, Beltran Rodriguez-Mueller, Yu Hui Rogers, Matthew C. Ross, Carsten Russ, Ravi K. Sanka, Pamela Sankar, J. Fah Sathirapongsasuti, Jeffery A. Schloss, Patrick D. Schloss, Thomas M. Schmidt, Matthew Scholz, Lynn Schriml, Alyxandria M. Schubert, Nicola Segata, Julia A. Segre, William D. Shannon, Richard R. Sharp, Thomas J. Sharpton, Narmada Shenoy, Nihar U. Sheth, Gina A. Simone, Indresh Singh, Christopher S. Smillie, Jack D. Sobel, Daniel D. Sommer, Paul Spicer, Granger G. Sutton, Sean M. Sykes, Diana G. Tabbaa, Mathangi Thiagarajan, Chad M. Tomlinson, Manolito Torralba, Todd J. Treangen, Rebecca M. Truty, Tatiana A. Vishnivetskaya, Jason Walker, Lu Wang, Zhengyuan Wang, Doyle V. Ward, Wesley Warren, Mark A. Watson, Christopher Wellington, Kris A. Wetterstrand, James R. White, Katarzyna Wilczek-Boney, Yuanqing Wu, Kristine M. Wylie, Todd Wylie, Chandri Yandava, Liang Ye, Yuzhen Ye, Shibu Yooseph, Bonnie P. Youmans, Lan Zhang, Yanjiao Zhou, Yiming Zhu, Laurie Zoloth, Jeremy D. Zucker, Bruce W. Birren, Richard A. Gibbs, Sarah K. Highlander, Barbara A. Methé, Karen E. Nelson, Joseph F. Petrosino, George M. Weinstock, Richard K. Wilson, and Owen White. Structure, function and diversity of the healthy human microbiome. Nature, 486(7402): 207–214, 2012. ISSN 00280836. doi: 10.1038/nature11234. URL http://dx.doi.org/10.1038/nature11234.

Eliran Kadosh, Irit Snir-Alkalay, Avanthika Venkatachalam, Shahaf May, Audrey Lasry, Ela Elyada, Adar Zinger, Maya Shaham, Gitit Vaalani, Marco Mernberger, and et al. The gut microbiome switches mutant p53 from tumour-suppressive to oncogenic. Nature, 586(7827): 133–138, 2020. doi: 10.1038/s41586-020-2541-0.

Dongwan Kang, Feng Li, Edward S Kirton, Ashleigh Thomas, Rob S Egan, Hong An, and Zhong Wang. Metabat 2: An adaptive binning algorithm for robust and efficient genome reconstruction from metagenome assemblies. PeerJ, 2019. doi: 10.7287/peerj.preprints.27522v1.

Fabian Kern, Tobias Fehlmann, and Andreas Keller. On the lifetime of bioinformatics web services. Nucleic Acids Research, 48(22): 12523–12533, December 2020. ISSN 0305-1048, 1362-4962. doi: 10.1093/nar/gkaa1125. URL https://academic.oup.com/nar/article/48/22/12523/6018434.

Kristopher Kieft, Zhichao Zhou, and Karthik Anantharaman. VIBRANT: Automated recovery, annotation and curation of microbial viruses, and evaluation of viral community function from genomic sequences. Microbiome, 8(1): 1–23, 2020. ISSN 20492618. doi: 10.1186/s40168-020-00867-0.

Silas Kieser, Joseph Brown, Evgeny M. Zdobnov, Mirko Trajkovski, and Lee Ann McCue. Atlas: A snakemake workflow for assembly, annotation, and genomic binning of metagenome sequence data. BMC Bioinformatics, 2019. doi: 10.1101/737528.

Mikhail Kolmogorov, Derek M. Bickhart, Bahar Behsaz, Alexey Gurevich, Mikhail Rayko, Sung Bong Shin, Kristen Kuhn, Jeffrey Yuan, Evgeny Polevikov, Timothy P.L. Smith, and Pavel A. Pevzner. metaFlye: scalable long-read metagenome assembly using repeat graphs. Nature Methods, 17(11): 1103–1110, 2020. ISSN 15487105. doi: 10.1038/s41592-020-00971-x. URL http://dx.doi.org/10.1038/s41592-020-00971-x.

Sabrina Krakau, Daniel Straub, Hadrien Gourlé, Gisela Gabernet, and Sven Nahnsen. Nf-core/mag: A best-practice pipeline for metagenome hybrid assembly and binning. NAR Genomics and Bioinformatics, 2021. doi: 10.1101/2021.08.29.458094.

Gregory M et. al. Kurtzer. Singularity 2.5.2 - linux application and environment containers for science, Jul 2018. URL https://zenodo.org/record/1308868.

Ben Langmead and Steven L. Salzberg. Fast gapped-read alignment with Bowtie 2. Nature Methods, 9(4): 357–359, 2012. ISSN 15487091. doi: 10.1038/nmeth.1923.

Dinghua Li, Chi Man Liu, Ruibang Luo, Kunihiko Sadakane, and Tak Wah Lam. MEGAHIT: An ultra-fast single-node solution for large and complex metagenomics assembly via succinct de Bruijn graph. Bioinformatics, 31(10): 1674–1676, 2015. ISSN 14602059. doi: 10.1093/bioinformatics/btv033.

Heng Li. Minimap2: Pairwise alignment for nucleotide sequences. Bioinformatics, 34(18): 3094–3100, 2018. ISSN 14602059. doi: 10.1093/bioinformatics/bty191.

Heng Li, Bob Handsaker, Alec Wysoker, Tim Fennell, Jue Ruan, Nils Homer, Gabor Marth, Goncalo Abecasis, and Richard Durbin. The Sequence Alignment/Map format and SAMtools. Bioinformatics, 25(16): 2078–2079, 2009. ISSN 13674803. doi: 10.1093/bioinformatics/btp352.

Oxford Nanopore Technologies Ltd. Medaka, 2019. URL https://github.com/nanoporetech/medaka.

Jennifer Lu, Florian P. Breitwieser, Peter Thielen, and Steven L. Salzberg. Bracken: Estimating species abundance in metagenomics data. PeerJ Computer Science, 3, 2017. doi: 10.7717/peerj-cs.104.

Serghei Mangul, Lana S. Martin, Eleazar Eskin, and Ran Blekhman. Improving the usability and archival stability of bioinformatics software. Genome Biology, 20(1), 2019. doi: 10.1186/s13059-019-1649-8.

Jose Manuel Martí. Recentrifuge: Robust comparative analysis and contamination removal for metagenomics. PLoS Computational Biology, 15(4): 1–24, 2019. ISSN 15537358. doi: 10.1371/journal.pcbi.1006967.

Guillaume Marçais, Arthur L. Delcher, Adam M. Phillippy, Rachel Coston, Steven L. Salzberg, and Aleksey Zimin. Mummer4: A fast and versatile genome alignment system. PLOS Computational Biology, 14(1):e1005944, January 2018. ISSN 1553-7358. doi: 10.1371/journal.pcbi.1005944. URL https://dx.plos.org/10.1371/journal.pcbi.1005944.

Dirk Merkel. Docker: lightweight linux containers for consistent development and deployment. Linux J., 2014(239): 2, 2014.

Cristina Moraru. VirClust—A Tool for Hierarchical Clustering, Core Protein Detection and Annotation of (Prokaryotic) Viruses. Viruses, 15(4), 2023. ISSN 19994915. doi: 10.3390/v15041007.

Eli L. Moss, Dylan G. Maghini, and Ami S. Bhatt. Complete, closed bacterial genomes from microbiomes using nanopore sequencing. Nature Biotechnology, 38(6): 701–707, 2020. doi: 10.1038/s41587-020-0422-6.

Felix Mölder, Kim Philipp Jablonski, Brice Letcher, Michael B. Hall, Christopher H. Tomkins-Tinch, Vanessa Sochat, Jan Forster, Soohyun Lee, Sven O. Twardziok, Alexander Kanitz, and et al. Sustainable data analysis with snakemake. F1000Research, 10:33, 2021. doi: 10.12688/f1000research.29032.1.

Stephen Nayfach, Antonio Pedro Camargo, Frederik Schulz, Emiley Eloe-Fadrosh, Simon Roux, and Nikos C. Kyrpides. CheckV assesses the quality and completeness of metagenome-assembled viral genomes. Nature Biotechnology, 39(5): 578–585, 2021. ISSN 15461696. doi: 10.1038/s41587-020-00774-7. URL http://dx.doi.org/10.1038/s41587-020-00774-7.

National Bioinformatics Infrastructure Sweden (NBIS). Nbis-meta: Snakemake workflow for metagenomic assembly, binning, and analysis, 2022. URL https://github.com/NBISweden/nbis-meta.

Sergey I Nikolenko, Anton I Korobeynikov, and Max A Alekseyev. Bayeshammer: Bayesian clustering for error correction in single-cell sequencing. BMC Genomics, 14(Suppl 1), 2013. doi: 10.1186/1471-2164-14-s1-s7.

Jakob Nybo Nissen, Joachim Johansen, Rosa Lundbye Allesøe, Casper Kaae Sønderby, Jose Juan Almagro Armenteros, Christopher Heje Grønbech, Lars Juhl Jensen, Henrik Bjørn Nielsen, Thomas Nordahl Petersen, Ole Winther, and Simon Rasmussen. Improved metagenome binning and assembly using deep variational autoencoders. Nature Biotechnology, 39(5):555– 560, may 2021. ISSN 15461696. doi: 10.1038/s41587-020-00777-4. URL https://doi.org/10.1038/s41587-020-00777-4.

Sergey Nurk, Dmitry Meleshko, Anton Korobeynikov, and Pavel A. Pevzner. MetaSPAdes: A new versatile metagenomic assembler. Genome Research, 27(5): 824–834, 2017. ISSN 15495469. doi: 10.1101/gr.213959.116.

Matthew R Olm, Christopher T Brown, Brandon Brooks, and Jillian F Banfield. drep: a tool for fast and accurate genomic comparisons that enables improved genome recovery from metagenomes through de-replication. The ISME Journal, 11(12): 2864–2868, Dec 2017. ISSN 1751-7362, 1751-7370. doi: 10.1038/ismej.2017.126. URL https://www.nature.com/articles/ismej2017126.

Askarbek Orakov, Anthony Fullam, Luis Pedro Coelho, Supriya Khedkar, Damian Szklarczyk, Daniel R. Mende, Thomas S.B. Schmidt, and Peer Bork. GUNC: detection of chimerism and contamination in prokaryotic genomes. Genome Biology, 22(1): 1–19, 2021. ISSN 1474760X. doi: 10.1186/s13059-021-02393-0.

Julie Orjuela, Aurore Comte, Sébastien Ravel, Florian Charriat, Tram Vi, François Sabot, and Sébastien Cunnac. Culebront: A streamlined long reads multi-assembler pipeline for prokaryotic and eukaryotic genomes. Peer Community Journal, 2, 2022. doi: 10.24072/pcjournal.153.

Shaojun Pan, Chengkai Zhu, Xing Ming Zhao, and Luis Pedro Coelho. A deep siamese neural network improves metagenome-assembled genomes in microbiome datasets across different environments. Nature Communications, 13(1): 1–12, 2022. ISSN 20411723. doi: 10.1038/s41467-022-29843-y.

Donovan H. Parks, Michael Imelfort, Connor T. Skennerton, Philip Hugenholtz, and Gene W. Tyson. Checkm: assessing the quality of microbial genomes recovered from isolates, single cells, and metagenomes. Genome Research, 25(7):1043–1055, July 2015. ISSN 1088-9051, 1549-5469. doi: 10.1101/gr.186072.114. URL http://genome.cshlp.org/lookup/doi/10.1101/gr.186072.114.

Donovan H. Parks, Christian Rinke, Maria Chuvochina, Pierre-Alain Chaumeil, Ben J. Woodcroft, Paul N. Evans, Philip Hugenholtz, and Gene W. Tyson. Recovery of nearly 8,000 metagenome-assembled genomes substantially expands the tree of life. Nature Microbiology, 2(11): 1533–1542, 2017. doi: 10.1038/s41564-017-0012-7. URL https://doi.org/10.1038/s41564-017-0012-7.

Vincent Prost, Stéphane Gazut, and Thomas Brüls. A zero inflated log-normal model for inference of sparse microbial association networks. PLOS Computational Biology, 17(6):e1009089, 2021.

Christopher Quince, Alan W Walker, Jared T Simpson, Nicholas J Loman, and Nicola Segata. Shotgun metagenomics, from sampling to analysis. Nature Biotechnology, 35(9): 833–844, 2017. doi: 10.1038/nbt.3935.

A Bramham Reddy and Karumathil P Gopinathan. Characterization of genomic dna of mycobacteriophage i3. Current Microbiology, 14: 241–245, 1987.

Jie Ren, Kai Song, Chao Deng, Nathan A. Ahlgren, Jed A. Fuhrman, Yi Li, Xiaohui Xie, Ryan Poplin, and Fengzhu Sun. Identifying viruses from metagenomic data using deep learning. Quantitative Biology, 8(1): 64–77, 2020. ISSN 20954697. doi: 10.1007/s40484-019-0187-4.

Alexa Ross, Samantha Ward, and Paul Hyman. More is better: selecting for broad host range bacteriophages. Frontiers in microbiology, 7:1352, 2016.

Hans-Joachim Ruscheweyh, Alessio Milanese, Lucas Paoli, Anna Sintsova, Daniel R Mende, Georg Zeller, and Shinichi Sunagawa. motus: profiling taxonomic composition, transcriptional activity and strain populations of microbial communities. Current Protocols, 1(8):e218, 2021.

Sara Saheb Kashaf, Alexandre Almeida, Julia A. Segre, and Robert D. Finn. Recovering prokaryotic genomes from host-associated, short-read shotgun metagenomic sequencing data. Nature Protocols, 16(5): 2520–2541, 2021. doi: 10.1038/s41596-021-00508-2.

Oliver Schwengers, Lukas Jelonek, Marius Alfred Dieckmann, Sebastian Beyvers, Jochen Blom, and Alexander Goesmann. Bakta: Rapid and standardized annotation of bacterial genomes via alignment-free sequence identification. Microbial Genomics, 7 (11), 2021. ISSN 20575858. doi: 10.1099/MGEN.0.000685.

Alexander Sczyrba, Peter Hofmann, Peter Belmann, David Koslicki, Stefan Janssen, Johannes Dröge, Ivan Gregor, Stephan Majda, Jessika Fiedler, Eik Dahms, and et al. Critical assessment of metagenome interpretation—a benchmark of metagenomics software. Nature Methods, 14(11): 1063–1071, 2017. doi: 10.1038/nmeth.4458.

Torsten Seemann. Prokka: Rapid prokaryotic genome annotation. Bioinformatics, 30(14): 2068–2069, 2014. ISSN 14602059. doi: 10.1093/bioinformatics/btu153.

Christian M. Sieber, Alexander J. Probst, Allison Sharrar, Brian C. Thomas, Matthias Hess, Susannah G. Tringe, and Jillian F. Banfield. Recovery of genomes from metagenomes via a dereplication, aggregation, and scoring strategy. Nature Microbiology, 2017. doi: 10.1101/107789.

Maria A. Sierra, Krista A. Ryon, Braden T. Tierney, Jonathan Foox, Chandrima Bhattacharya, Evan Afshin, Daniel Butler, Stefan J. Green, W. Kelley Thomas, Jordan Ramsdell, and et al. Microbiome and metagenomic analysis of lake hillier australia reveals pigment-rich polyextremophiles and wide-ranging metabolic adaptations. Environmental Microbiome, 17(1), 2022. doi: 10.1186/s40793-022-00455-9.

Renaud Van Damme, Martin Hölzer, Adrian Viehweger, Bettina Müller, Erik Bongcam-Rudloff, and Christian Brandt. Metagenomics workflow for hybrid assembly, differential coverage binning, metatranscriptomics and pathway analysis (muffin). PLOS Computational Biology, 17(2), 2021. doi: 10.1371/journal.pcbi.1008716.

Riccardo Vicedomini, Christopher Quince, Aaron E. Darling, and Rayan Chikhi. Strainberry: Automated strain separation in low-complexity metagenomes using long reads. Nature Communications, 12(1), 2021. doi: 10.1038/s41467-021-24515-9.

Ziye Wang, Pingqin Huang, Ronghui You, Fengzhu Sun, and Shanfeng Zhu. MetaBinner: a high-performance and stand-alone ensemble binning method to recover individual genomes from complex microbial communities. Genome Biology, 24(1): 1–18, 2023. ISSN 1474760X. doi: 10.1186/s13059-022-02832-6. URL https://doi.org/10.1186/s13059-022-02832-6.

Ryan R. Wick. Porechop, 2018. URL https://github.com/rrwick/Porechop.

Timothy J. Williams, Michelle A. Allen, Angelique E. Ray, Nicole Benaud, Devan S. Chelliah, Davide Albanese, Claudio Donati, Laura Selbmann, Claudia Coleine, and Belinda C. Ferrari. Novel endolithic bacteria of phylum chloroflexota reveal a myriad of potential survival strategies in the antarctic desert. Applied and Environmental Microbiology, pages e02264–23, February 2024. ISSN 0099-2240, 1098–5336. doi: 10.1128/aem.02264-23. URL https://journals.asm.org/doi/10.1128/aem.02264-23.

Derrick E. Wood, Jennifer Lu, and Ben Langmead. Improved metagenomic analysis with kraken 2. Genome Biology, 2019. doi: 10.1101/762302.

Yu Wei Wu, Blake A. Simmons, and Steven W. Singer. MaxBin 2.0: An automated binning algorithm to recover genomes from multiple metagenomic datasets. Bioinformatics, 32(4):605–607, feb 2016. ISSN 14602059. doi: 10.1093/bioinformatics/btv638. URL https://www.jbei.org.

